# Identification of overlapping and distinct mural cell populations during early embryonic development

**DOI:** 10.1101/2023.04.03.535476

**Authors:** Sarah Colijn, Miku Nambara, Amber N. Stratman

**Affiliations:** Department of Cell Biology and Physiology, Washington University in St. Louis, School of Medicine, St. Louis, MO 63110

**Keywords:** mural cell progenitor, vSMC, pericyte, perivascular cells, hypochord, sclerotome, zebrafish

## Abstract

Mural cells are an essential perivascular cell population that associate with blood vessels and contribute to vascular stabilization and tone. In the embryonic zebrafish vasculature, *pdgfrb* and *tagln* are commonly used as markers for identifying pericytes and vascular smooth muscle cells (vSMCs). However, the expression patterns of these markers used in tandem have not been fully described. Here, we used the *Tg(pdgfrb:Gal4FF; UAS:RFP)* and *Tg(tagln:NLS-EGFP)* transgenic lines to identify single- and double-positive perivascular populations in the cranial, axial, and intersegmental vessels between 1 and 5 days post-fertilization. From this comparative analysis, we discovered two novel regions of *tagln*-positive cell populations that have the potential to function as mural cell precursors. Specifically, we found that the hypochord— a reportedly transient structure—contributes to *tagln*-positive cells along the dorsal aorta. We also identified a unique sclerotome-derived mural cell progenitor population that resides along the midline between the neural tube and notochord and contributes to intersegmental vessel mural cell coverage. Together, our findings highlight the variability and versatility of tracking *pdgfrb* and *tagln* expression in mural cells of the developing zebrafish embryo.

**Summary Statement:** Detailed analysis of *pdgfrb*/*tagln* vascular expression patterns in embryonic zebrafish reveals novel regions of *tagln*-positive populations such as the hypochord and a mural cell progenitor population adjacent to the midline.

## Introduction

In vertebrate development, early blood vessels are initially comprised of a single layer of endothelial cells—which then recruit mural cells to assist in basement membrane deposition, vessel contractility, and cellular communication (Armulik et al., 2005). Mural cells can vary widely in morphology and gene expression patterns depending on vessel type, vascular bed, tissue origin, differentiation status, developmental stage, and more (Donadon and Santoro, 2021, Owens et al., 2004, Holm et al., 2018). Types of mural cells include pericytes—which are associated with smaller vessels such as capillaries—and vascular smooth muscle cells (vSMCs) on larger vessels. vSMCs will accumulate on arteries and—to a lesser extent—on veins during vascular development to provide tensile strength and promote basement membrane deposition (Stratman et al., 2017). Due to the diverse properties of mural cells, different gene expression profiles can be used to identify mural cell types along various regions of the vasculature. Therefore, identification of appropriate cellular markers for distinct developmental stages is necessary for the study of vessel stabilization in different vascular beds.

In the embryonic zebrafish, platelet-derived growth factor receptor beta (*pdgfrb*) and transgelin (*tagln*) are commonly used as markers to identify pericytes and vSMCs. However, the temporal expression patterns of these markers used in tandem has not been fully described. Pdgfrb—a surface receptor for the chemoattractant ligands of the Pdgf family—is used as a marker for pericytes and immature vSMCs (Hellstrom et al., 1999, Armulik et al., 2005, Ando et al., 2021b). Pdgfrb has been shown to be required for mural cell expansion both embryonically and postnatally (Leonard et al., 2022, Ando et al., 2021b, Stratman et al., 2020). Cranial and cardiac neural crest cells also express Pdgfrb and have the capacity to become mural cells in addition to giving rise to other mesenchymal cell populations (Smith and Tallquist, 2010, Majesky, 2007, Ando et al., 2016). Some mature mural cells—such as pericytes—continue to express Pdgfrb, though less than 1% of Pdgfrb-positive cells in a zebrafish larvae at 5 days post-fertilization (dpf) are pericytes (Shih et al., 2021). This suggests that most Pdgfrb-positive cells in early development are progenitors for diverse cell populations.

Tagln (or SM22)—an actin-binding protein from the calponin family—is commonly used to identify mature, contractile vSMCs (Chakraborty et al., 2019, Santoro et al., 2009). Tagln and other actin-binding proteins confer contractile function to mature mural cells, allowing them to regulate vascular tone and blood pressure (Rensen et al., 2007). During development, Tagln is also expressed in intestinal smooth muscle, cardiac muscle, and skeletal muscle cells (Seiler et al., 2010, Li et al., 1996), though these muscle cells can be distinguished from vSMCs by observing cellular morphology and proximity to the vasculature. Altogether, this suggests that the *pdgfrb* and *tagln* reporters can be used to identify mural cells and their differentiation status by tracking single- versus double-positive cells in discrete vascular beds over the course of early development.

In this report, we use *Tg(pdgfrb:Gal4FF; UAS:RFP)* and *Tg(tagln:NLS-EGFP)* transgenic lines to identify single- and double-positive pericytes/vSMCs in the cranial, axial, and intersegmental vessels (ISVs) at 1 to 5 dpf. Through this study, we have uncovered two novel sites of *tagln*-positive cell populations that have the potential to function as mural cell precursors in the zebrafish trunk. First, we found that the hypochord—a reportedly transient structure present in fish and amphibians—contributes to *tagln*-positive cells along the dorsal aorta. Second, we identified a sclerotome-derived mural cell progenitor population that resides along the midline at the notochord-neural tube interface and contributes to ISV mural cell coverage. These findings highlight unexpected and nuanced gene expression patterns across perivascular cell maturation.

## Results

### *pdgfrb* and *tagln* transgenic reporter lines

To visualize mural cells in the zebrafish embryo, we crossed the established transgenic lines *TgBAC(pdgfrb:Gal4FF)^ncv24^, Tg(5xUAS:RFP)^zf83^,* and *Tg(tagln:NLS-EGFP-2A-CFP-FTASE)^y450^*to generate embryos that were heterozygous carriers for each transgene. Expression of both *pdgfrb* and *tagln* were notable at all embryonic stages analyzed—with the brightest signal for *pdgfrb* visible in the anterior and cranial regions, and *tagln* in the skeletal muscle of the trunk and later in the intestinal compartment (Fig. S1).

The *pdgfrb* reporter line utilizes the Gal4/UAS system (Ando et al., 2016)— designated as *Tg(pdgfrb:Gal4FF; UAS:RFP)* in this report—and mosaic fluorescence was occasionally observed during screening; embryos with the least mosaicism were selected for use. The *tagln* reporter line—designated as *Tg(tagln:NLS-EGFP)—*contains a nuclear localization signal that allows for GFP to be concentrated in the nucleus (Stratman et al., 2017), a modification that facilitates the identification of individual *tagln*-positive cells in regions with dense *tagln* expression. With this tool, our study provides a unique approach to tracking *pdgfrb* and *tagln* localization in the zebrafish.

### Imaging the mural cells of the cranial vasculature

Development of the zebrafish cranial vasculature has been thoroughly mapped through numerous studies. This process involves the formation of the primordial hind- and midbrain channels (PHBC/PMBC) and the lateral dorsal aorta (LDA) by 1 dpf, the growth of the middle cerebral vein (MCeV), metencephalic artery (MtA), basilar artery (BA), and Circle of Willis (CoW) by 1.5 dpf, and the sprouting and fusion of various central arteries (CtAs) from the PHBC to the BA by 2 dpf (Ulrich et al., 2011, Fujita et al., 2011, Isogai et al., 2001). These vessels continue to remodel, grow, and stabilize, though the patterning remains mostly intact from 6 dpf and beyond (Isogai et al., 2001).

Mural cell coverage of the cranial vessels has also been frequently studied, with the common understanding being that neural crest-derived *pdgfrb*-positive pericytes are the predominant mural cell in the mid- and hindbrain prior to 5 dpf (Ando et al., 2019, Ando et al., 2016, Ando et al., 2021a, Wang et al., 2014). Interestingly, Ando et al have shown that some mural cells in the hindbrain are derived from mesoderm. Specifically, *pdgfrb*-positive cells surrounding the PHBC at 1 dpf are purportedly mesoderm-derived mesenchyme; these cells then increase *pdgfrb* expression, migrate to CtAs, and eventually acquire vSMC markers—such as *tagln*—at 5 dpf (Ando et al., 2016, Ando et al., 2021a). Prior to 5 dpf, vSMC markers are sparse in the cranial vessels, with less than 10% of mural cells on the CtAs expressing *tagln* by 5 dpf (Ando et al., 2019).

In our *Tg(tagln:NLS-EGFP); Tg(pdgfrb:Gal4FF; UAS:RFP)* embryos, as expected, *pdgfrb* was the predominant marker of cranial mural cells between 1 and 5 dpf (Figs 1A-J; S2A-K). At 1 dpf, *pdgfrb*-positive cells were loosely associated with the PHBC (Fig. 1B)—corroborating previous data (Ando et al., 2021a, Ando et al., 2016)—and with the MtA and MCeV (Fig. S2B,C). Based on the current understanding of the literature, these early *pdgfrb*-positive cells are likely a combination of neural crest and mesoderm-derived mesenchyme that have the potential to migrate to the vessels and continue to mature into mural cells throughout development.

**Figure 1:**
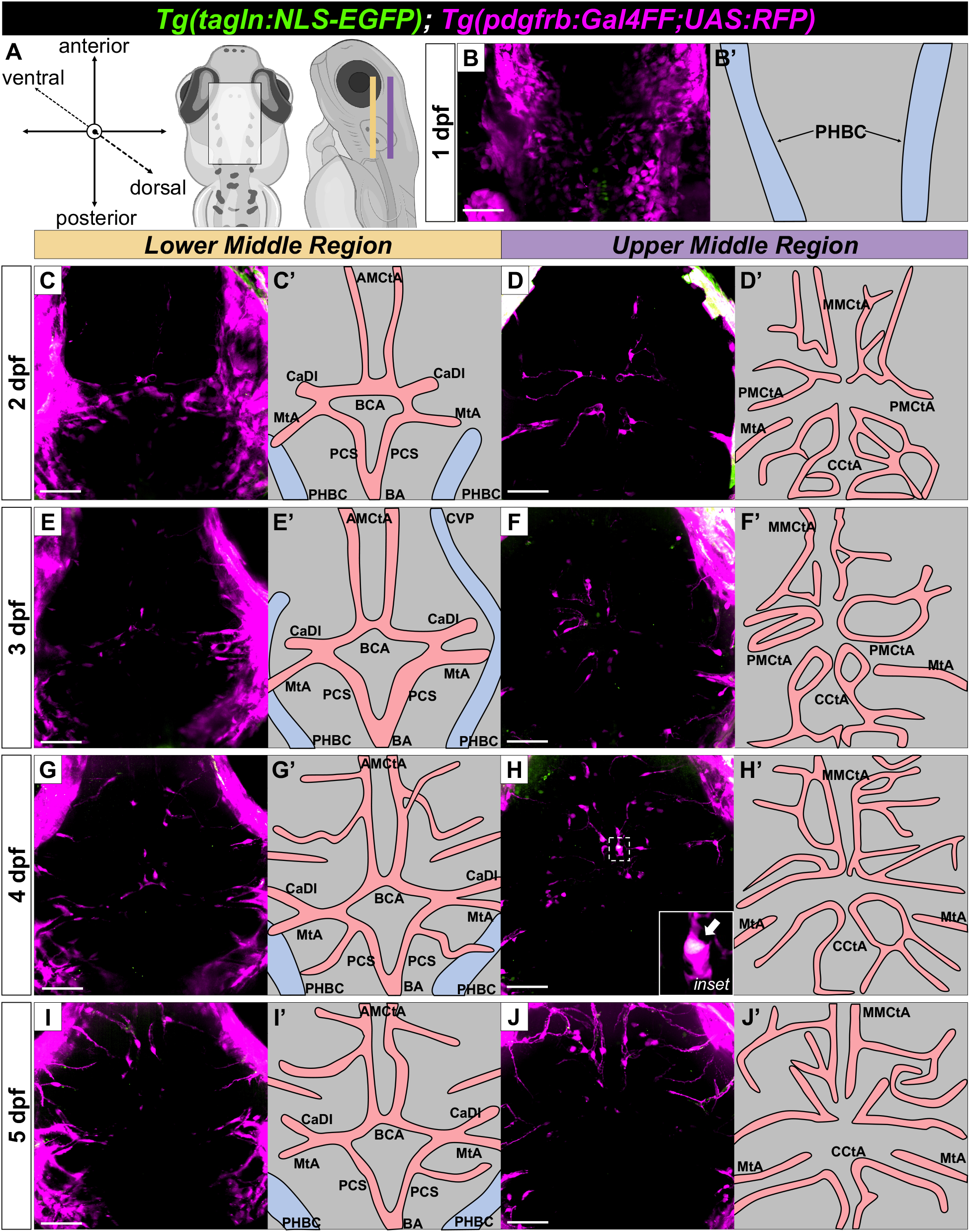
Coronal views of *Tg(tagln:NLS-EGFP); Tg(pdgfrb:Gal4FF; UAS:RFP)* zebrafish brains from 1 to 5 dpf. (A) Schematic diagram illustrating the orientation for imaging brain coronal sections: the lower middle region (yellow) and upper middle region (purple). (B-J) Maximum intensity projection images of the mid- and hindbrain from 1 to 5 dpf. (B’-J’) Outlines of the blood vessels visualized in the confocal images; pink represents arteries and blue represents veins. At 1 dpf (B) *pdgfrb*-positive cells are associated with the PHBC. At 2 dpf (C,D) *pdgfrb*-positive cells begin to accumulate around the CoW (C, comprised of the BCA and PCS) and on the CtAs (D). *pdgfrb*-positive cells continue to accumulate on the CoW and CtAs from 3 to 5 dpf (E-J). A single *pdgfrb*/*tagln* double-positive cell is evident on the MMCtA at 4 dpf (H, inset). Schematic created with BioRender.com. AMCtA, anterior mesencephalic central artery. BA, basilar artery. BCA, basal communicating artery. CaDI, caudal division of the internal carotid artery. CCtA, cerebellar central artery. CoW, Circle of Willis. MMCtA, middle mesencephalic central artery. MtA, metencephalic artery. PCS, posterior communicating segment. PHBC, primordial hindbrain channel. PMCtA, posterior mesencephalic central artery. Scale bars: 50 µm.

After 1 dpf, *pdgfrb*-positive cells slowly accumulate on the CoW and CtAs until 5 dpf, the latest stage analyzed (Figs 1C-J; S2D-K). A single *pdgfrb*/*tagln* double-positive cell can be seen at 4 dpf on the middle mesencephalic central artery (MMCtA; Fig. 1H, inset). However, throughout this region, *tagln*-positive cells were extremely sparse at all developmental stages assessed, consistent with previous reports that vSMC markers such as *tagln* are not prevalent in cranial mural cells prior to 4 dpf. These results indicate that *pdgfrb* is the predominant mural cell marker along embryonic cranial vessels, though a small proportion of these cells—particularly on the arteries—can mature into *pdgfrb*/*tagln* double-positive vSMCs by 5 dpf.

### Imaging the mural cells of the axial vasculature

The main axial vessels—i.e. dorsal aorta (DA) and posterior cardinal vein (PCV) in the anterior trunk and the caudal artery (CA) and caudal vein (CV) in the tail—are some of the first vessels to form prior to the onset of circulation at 1 dpf (Isogai et al., 2001). The mural cells that accumulate on the ventral side of the DA have been well-described; *pdgfrb*-positive cells begin to accumulate at 1.5 dpf and then acquire the vSMC marker *tagln* at 3 dpf (Ando et al., 2021a, Stratman et al., 2017, Ando et al., 2016). Approximately 25% of vSMCs on the ventral side of the DA are reported to be *pdgfrb*/*tagln* double-positive by 5 dpf (Ando et al., 2019).

The mural cells that cover the dorsal side of the DA are not as well-defined. Intriguingly, the mural cells that accumulate on the dorsal and ventral sides of the DA are initially derived from different mesodermal origins in mice (Wasteson et al., 2008). This lineage difference has not been definitively shown in zebrafish. However, there are clear distinctions between the cells that initially populate the dorsal versus ventral sides of the DA in zebrafish embryos—with the mural cells on the dorsal side of the DA reported to express *tagln* but not *pdgfrb* (Ando et al., 2016).

By tracking the expression of *pdgfrb* and *tagln* in vSMCs on the DA, PCV, CA, and CV, we identify the hypochord as a novel source of *tagln*-positive cells on the dorsal side of the DA and CA (Fig. 2). We also corroborate the previous findings regarding vSMC expression patterns on the ventral side of the DA, and observe a slow accumulation of *pdgfrb* single-positive cells on the PCV and CV between 1 to 5 dpf (Fig. 2). Finally, we find that accumulation and maturation of these mural cells occurs in an anterior to posterior fashion, with mural cells on the anterior DA and PCV maturing one to two days earlier than the mural cells of the posterior CA and CV (Fig. 2).

**Figure 2:**
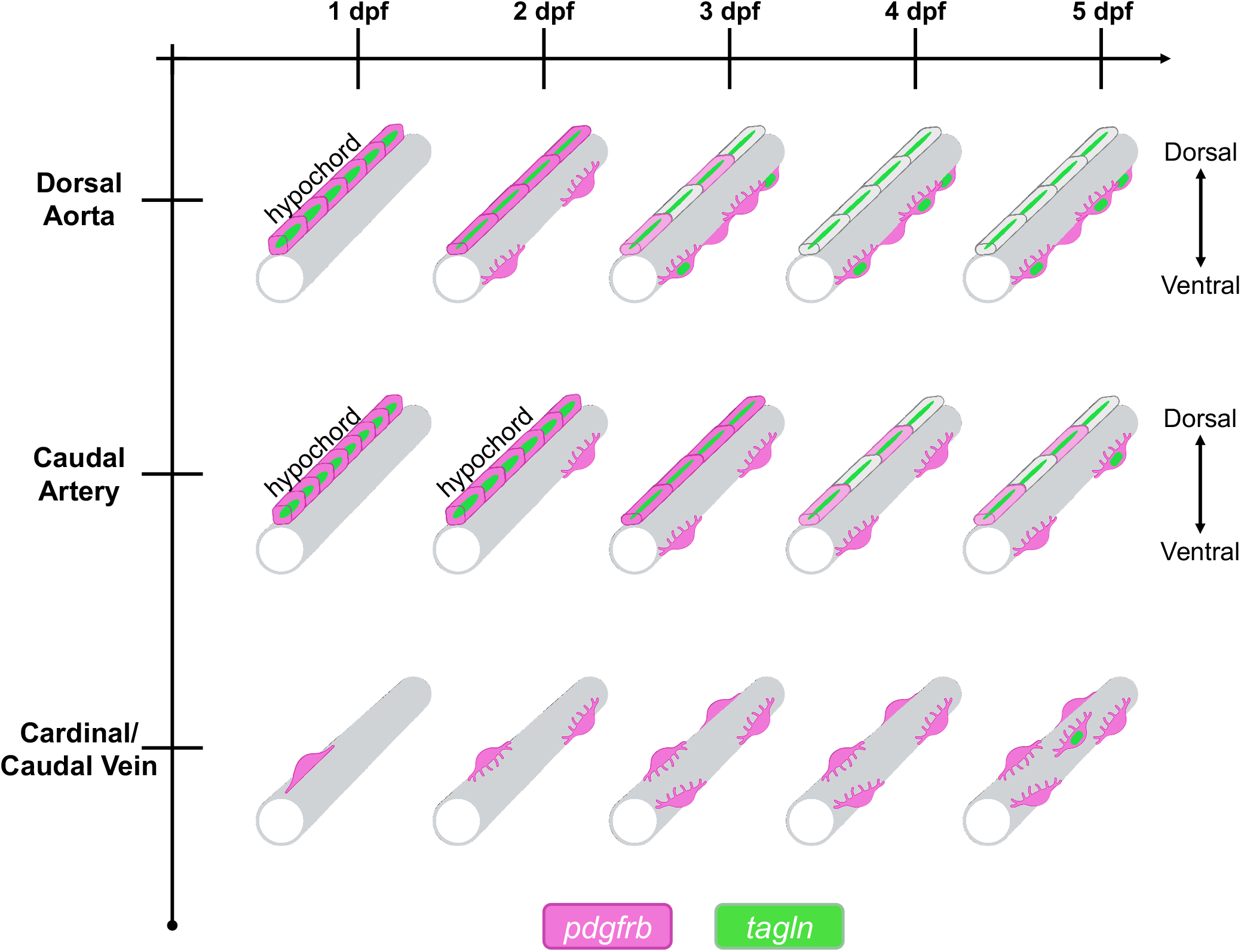
Summary of mural cell accumulation on the axial vasculature. The hypochord—which begins as a *pdgfrb*/*tagln* double-positive structure—is a novel source of *tagln*-positive cells on the dorsal side of the dorsal aorta and caudal artery. At 2 dpf, the hypochord cells began to delaminate on the dorsal aorta. At 3 dpf, the *pdgfrb*/*tagln* double-positive cells begin to decrease *pdgfrb* expression. By 4 dpf, these cells were *tagln* single-positive. In the caudal region, this hypochord reprogramming event began 1 day later than the anterior region. On the ventral side of the dorsal aorta and caudal vein, *pdgfrb* single-positive vSMCs were observed beginning at 2 dpf, where they then expanded in number and increased expression of *tagln*. By 5 dpf, *pdgfrb*/*tagln* double-positive cells accounted for approximately 60% of ventral vSMCs on the dorsal aorta, and no *tagln* single-positive cells were detected. The vSMCs on the ventral side of the CA accumulated more slowly and did not begin co-expressing *tagln* until 5 dpf, which amounted to a 2-day delay compared to the anterior region. Finally, we observed a slow accumulation of *pdgfrb* single-positive cells on the posterior cardinal vein and caudal vein between 1 to 5 dpf. *tagln*-positive cells were particularly sparse on the veins, but visible at later stages.

To begin, we analyzed *pdgfrb* and *tagln* expression in mural cells of the anterior DA and PCV in *Tg(tagln:NLS-EGFP); Tg(pdgfrb:Gal4FF; UAS:RFP)* embryos between 1 to 5 dpf (Fig. 3A-I). We identified the hypochord at 1 dpf as the *pdgfrb*-positive structure immediately dorsal to the DA that runs along the midline (Fig. 3A). The hypochord is a reportedly transient, rod-like structure—only one cell in width and height—that resides beneath the notochord and assists in the formation and patterning of the DA (Eriksson and Löfberg, 2000, Cleaver et al., 2000, Löfberg and Collazo, 1997). Once the DA is formed, the hypochord cells are reported to delaminate and degenerate—in part due to apoptosis (Löfberg and Collazo, 1997, Cleaver et al., 2000)—then by 2.5 dpf either disappear or become indistinguishable from the cells that surround the DA in the zebrafish embryo (Eriksson and Löfberg, 2000).

**Figure 3:**
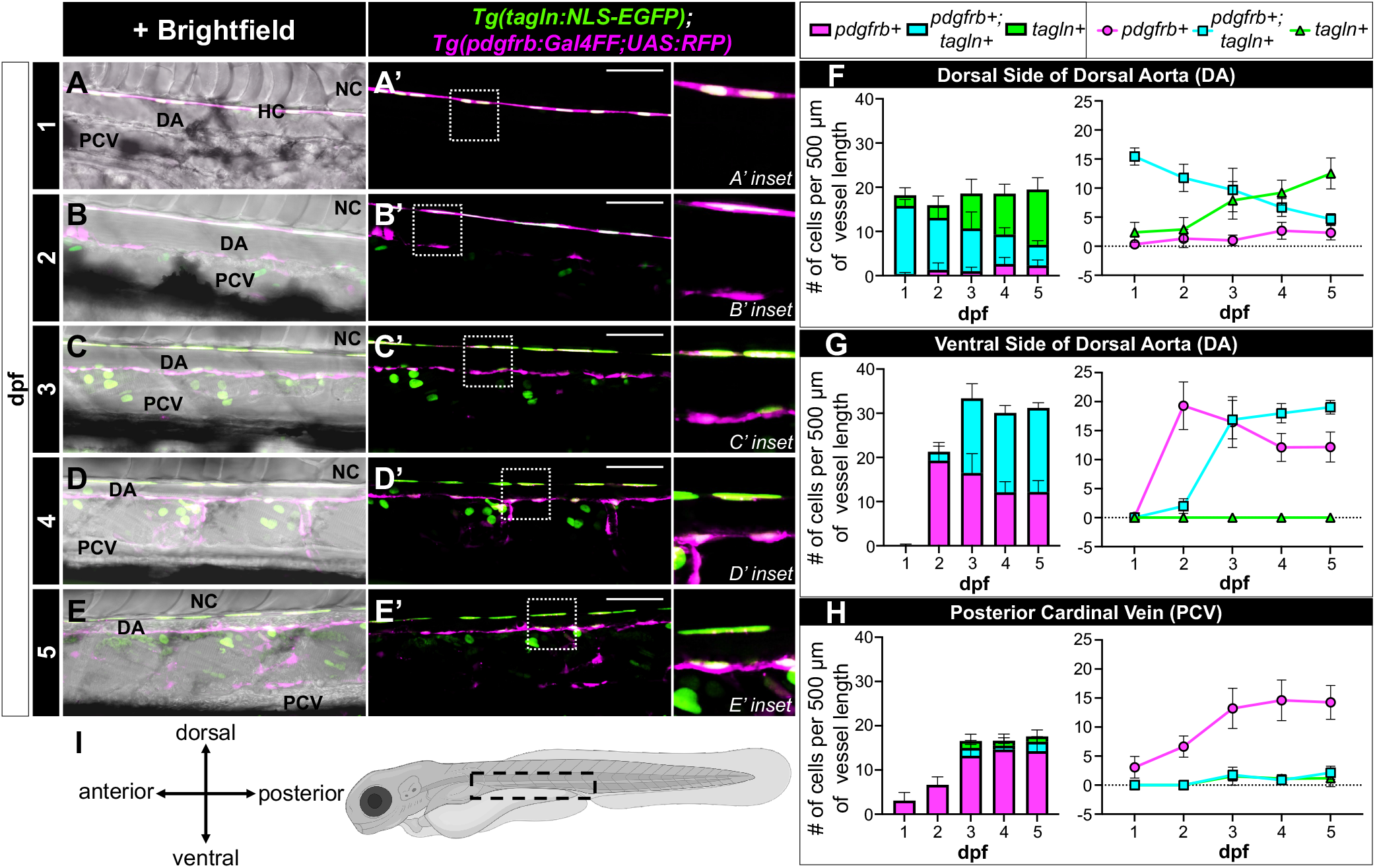
*tagln* and *pdgfrb* expression along the dorsal aorta and cardinal vein of 1 to 5 dpf *Tg(tagln:NLS-EGFP); Tg(pdgfrb:Gal4FF; UAS:RFP)* zebrafish. (A-E) Maximum intensity projection images of the DA and PCV from 1 to 5 dpf. (A’-E’) The images from A-E without the brightfield overlay. (F-G) Bar and line graph representations of the number of single- and double-positive cells on the dorsal side of the DA (F), the ventral side of the DA (G), and the PCV (H). (I) Schematic diagram illustrating the imaged region. Schematic created with BioRender.com. At least 6 embryos were analyzed at each age. DA, dorsal aorta. PCV, posterior cardinal vein. HC, hypochord. NC, notochord. Scale bars: 50 µm.

*pdgfrb* expression in the hypochord has been previously reported, in addition to *pdgfrb* expression in the floor plate—the structure that sits on the ventral side of the neural tube (Ando et al., 2019, Ando et al., 2016, Wiens et al., 2010). We found that in addition to *pdgfrb*, the hypochord also expresses *tagln* at the 1 dpf stage (Fig. 3A,F). At 2 dpf, the hypochord cells began to delaminate, as evidenced by the elongating nuclei (Fig. 3B). At 3 dpf, when the hypochord is allegedly absent from the anterior trunk (Eriksson and Löfberg, 2000), we observed that the *tagln*-positive cells persist and begin to decrease *pdgfrb* expression (Fig. 3C,F). By 4 dpf, these cells were *tagln* single-positive, with noticeably fewer nuclei visible per vessel length (Fig. 3D,F; Movie 1), indicating that a portion of hypochord cells do degenerate after delamination. However, a portion of elongated *tagln*-positive cells were still present at 5 dpf (Fig. 3E,F), and Ando et al reports observing these *tagln* single-positive cells above the DA as late as 7 dpf (Ando et al., 2016).

To confirm that these hypochord-derived cells are maintained—and that they are not ventral vSMCs that have migrated to the dorsal side of the DA—we used a sclerotome-marker twist family bHLH transcription factor 1a (*twist1a*) transgenic line to distinguish their origin (Stratman et al., 2017). The hypochord—in addition to the notochord and floor plate—has been shown to be derived from the multipotent midline progenitor cells in the dorsal organizer or the tailbud (Row et al., 2016, Latimer and Appel, 2006). Thus, hypochord cells maintain distinct origins from sclerotome cells, which emerge from the somites (Tani et al., 2020). Indeed, we found that the *tagln* single-positive cells on the dorsal side of the DA are not sclerotome-derived, unlike the vSMCs on the ventral side of the DA (Fig. S3A-F). Altogether, these results suggest that the hypochord is the source of the *tagln* single-positive cells that reside on the dorsal side of the DA. This also indicates that the cells of the hypochord are not entirely transient as previously assumed, but instead a portion reprogram their cellular function and morphology after the DA has been appropriately oriented.

We next observed the development of the mural cells on the ventral side of the DA. While the hypochord-derived cells on the dorsal side of the DA flattened and elongated over time, the vSMCs on the ventral side were more globular (Fig. 3A-E). *pdgfrb* single-positive vSMCs were observed beginning at 2 dpf, where they then expanded in number and increased expression of *tagln* (Fig. 3A-E,G). By 5 dpf, *pdgfrb*/*tagln* double-positive cells accounted for approximately 60% of ventral vSMCs, and no *tagln* single-positive cells were detected. These data are consistent with previous reports of vSMCs on the ventral side of the DA, though we observe a higher percentage of double-positive vSMCs (60% versus 25%) by 5 dpf (Ando et al., 2019, Ando et al., 2016, Ando et al., 2021a).

Furthermore, we detected *pdgfrb* single-positive cells associated with the PCV as early as 1 dpf—though these cells were sparse—followed by the slow and sporadic accumulation of *pdgfrb*-positive cells until 5 dpf (Fig. 3A-E,H). This timing is earlier than suggested in previous reports where *pdgfrb* expression was undetectable in cells around the PCV prior to 7 dpf (Ando et al., 2016). Stratman et al describes a small number of *tagln*-positive cells on the PCV around 4 dpf (Stratman et al., 2017). Similarly, we saw low numbers of weak *tagln*-positive cells beginning at 3 dpf, though these cells were less common than *pdgfrb* single-positive cells (Fig. 3H). These data indicate that mural cells of the PCV associate and mature at much later developmental stages than aortic vSMCs.

In the posterior trunk region, mural cells of the CA and CV associate with vessels and pattern similarly to the anterior mural cells but on a delayed time course (Fig. 4A-I). At 1 dpf, the hypochord cells were block-like and were still distinguishable as hypochord cells by 2 dpf (Fig. 4A,B,F). This was expected, as the hypochord “disappears” in an anterior to posterior fashion (Eriksson and Löfberg, 2000), and is evident at 2 dpf in the posterior region even when it is no longer detectable in the anterior region. We saw the hypochord cells had begun to delaminate by 3 dpf (Fig. 4C), at which point they appeared comparable to the 2 dpf hypochord cells in the anterior region (Fig. 3B). While the cells then began turning off *pdgfrb*, only about a third of hypochord-derived cells in the posterior trunk were *tagln* single-positive by 5 dpf (Fig. 4C-F), indicating a delay in hypochord reprogramming compared to the anterior hypochord.

**Figure 4:**
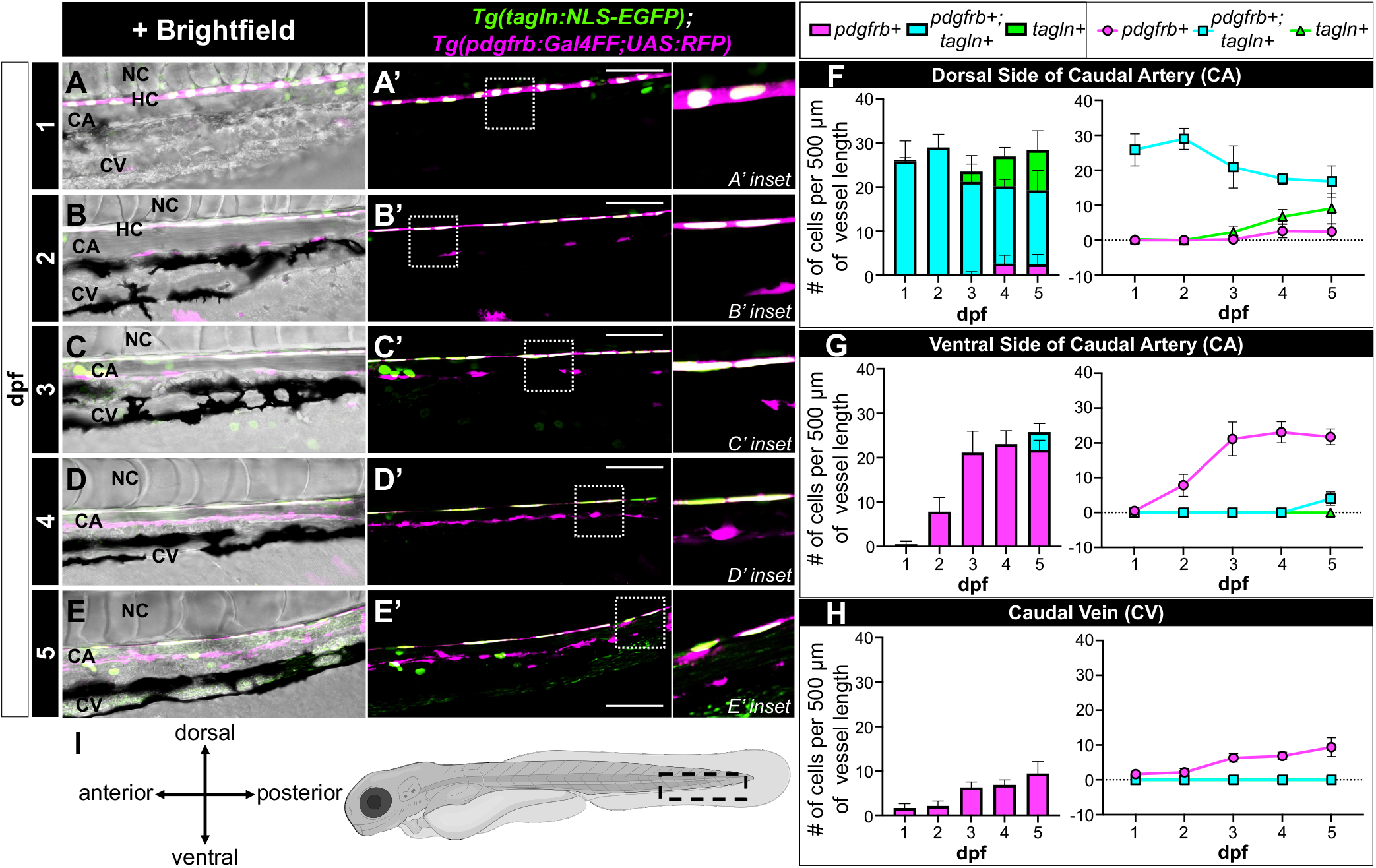
*tagln* and *pdgfrb* expression along the caudal artery and vein of 1 to 5 dpf *Tg(tagln:NLS-EGFP); Tg(pdgfrb:Gal4FF; UAS:RFP)* zebrafish. (A-E) Maximum intensity projection images of the CA and CV from 1 to 5 dpf. (A’-E’) The images from A-E without the brightfield overlay. (F-G) Bar and line graph representations of the number of single- and double-positive cells on the dorsal side of the CA (F), the ventral side of the CA (G), and the CV (H). (I) Schematic diagram illustrating the imaged region. Schematic created with BioRender.com. At least 6 embryos were analyzed at each age. CA, caudal artery. CV, caudal vein. HC, hypochord. NC, notochord. Scale bars: 50 µm.

Furthermore, the ventral side of the CA began acquiring *pdgfrb* single-positive cells at 2 dpf—similar to the DA—but they did not begin co-expressing *tagln* until 5 dpf (Fig. 4B-E,G). This amounted to a 2-day delay in mural cell development compared to the anterior region (Fig. 3G). Finally, mural cell coverage of the CV was particularly sparse (Fig. 4A-E,H), with half as many cells associated with the CV compared to the PCV at the same developmental stage (Fig. 3H). These data indicate that association and maturation of vSMCs on the main axial vessels occurs in an anterior to posterior fashion across time.

Of note, weak *tagln* expression was detected in endothelial cells of the axial vasculature (Fig. S4A,B), which is consistent with previous reports that the endothelium can express low levels of *tagln* (Chakraborty et al., 2019, Tsuji-Tamura et al., 2021). Despite this, *tagln* expression in the endothelial cells was easily distinguishable from vSMCs; therefore, it did not confound the mural cell counts of the axial vessels.

### Imaging the mural cells of the intersegmental vessels (ISVs)

We next analyzed the mural cell populations surrounding the ISVs of the trunk region. At 1 dpf, arterial ISVs (aISVs) begin to sprout from the DA and grow dorsally between the somites and the notochord/neural tube. At 1.5 dpf, venous ISVs (vISVs) sprout from the PCV, grow dorsally, and then with aISVs branch laterally to establish the dorsal longitudinal anastomotic vessel (DLAV) that resides above the neural tube (Isogai et al., 2003). By 2 dpf, the arterial and venous ISVs are mostly lumenized and reach from the DA/PCV below the notochord to the DLAV above the neural tube (Isogai et al., 2001). Mural cells of the ISVs have been reported to emerge by either *de novo* differentiation of nearby mesenchyme followed by subsequent proliferation or through migration from the ventral side of the DA (Ando et al., 2016). Studies identifying the origin of ISV mural cells show that they are primarily derived from the sclerotome (Stratman et al., 2017, Rajan et al., 2020)—a sub-compartment of the somites and a derivative of paraxial mesoderm (Tani et al., 2020). Zebrafish have recently been shown to have a dorsal and ventral sclerotome, both of which contribute to the cells that associate with the ISVs: dorsal sclerotome populates the ISV region adjacent to the neural tube while ventral sclerotome populates the full ISV (Rajan et al., 2020, Ma et al., 2018). However, while the dorsal and ventral sclerotome cells can contribute to the development of different regions of the embryo (Ma et al., 2018), it is currently unclear whether they differ in gene expression.

Rajan et al recently described a population of sclerotome-derived cells called perivascular fibroblasts that associate with ISVs by 2 dpf and produce collagen matrix, thus providing structural support prior to mural cell migration or differentiation (Rajan et al., 2020). These cells are primarily derived from ventral sclerotome, are evenly distributed between arterial and venous ISVs, remain constant in number on the ISVs from 2 to 4 dpf, and express low levels of *pdgfrb* (Rajan et al., 2020). At 3 dpf, a subset— but not all—of perivascular fibroblasts will increase *pdgfrb* expression to become ISV mural cells (Rajan et al., 2020). Ando et al also describes a naïve *pdgfrb*-low mesenchymal population—likely the same cell type or similar to perivascular fibroblasts— that can mature into *pdgfrb*-high mural cells on the ISVs starting at approximately 2.5 dpf (Ando et al., 2016, Ando et al., 2021a).

Both studies observe that mural cells with *pdgfrb*-high expression are more apparent on aISVs than vISVs (Ando et al., 2016, Rajan et al., 2020), which is in part due to maturation of *pdgfrb*-low mesenchyme to *pdgfrb*-high mural cells through Notch signaling in arterial endothelium (Ando et al., 2019). Ando et al also observes that approximately 15% of mural cells are *tagln*-positive on ISVs by 5 dpf (Ando et al., 2019), potentially indicating further maturation of these *pdgfrb*-positive cells.

In support of these previous studies, we observed *pdgfrb*-low and -high cell populations surrounding the ISVs up to 5 dpf in *Tg(tagln:NLS-EGFP); Tg(pdgfrb:Gal4FF; UAS:RFP)* embryos (Figs 5A-H; S5A-I). Unexpectedly, we also detected *tagln*-low and - high populations as early as 2 dpf (Figs 5C,D; S5A,B), prior to the developmental stages in which ISV mural cells reportedly mature. While a subset of *tagln*-low cells in this region were endothelial cells, we confirmed that many of the *tagln*-low and -high cells associated with the early ISVs were perivascular (Fig. S4C,D), indicating that *tagln* may not exclusively label mature mural cells, but instead also labels a subset of mural cell precursors.

**Figure 5.**
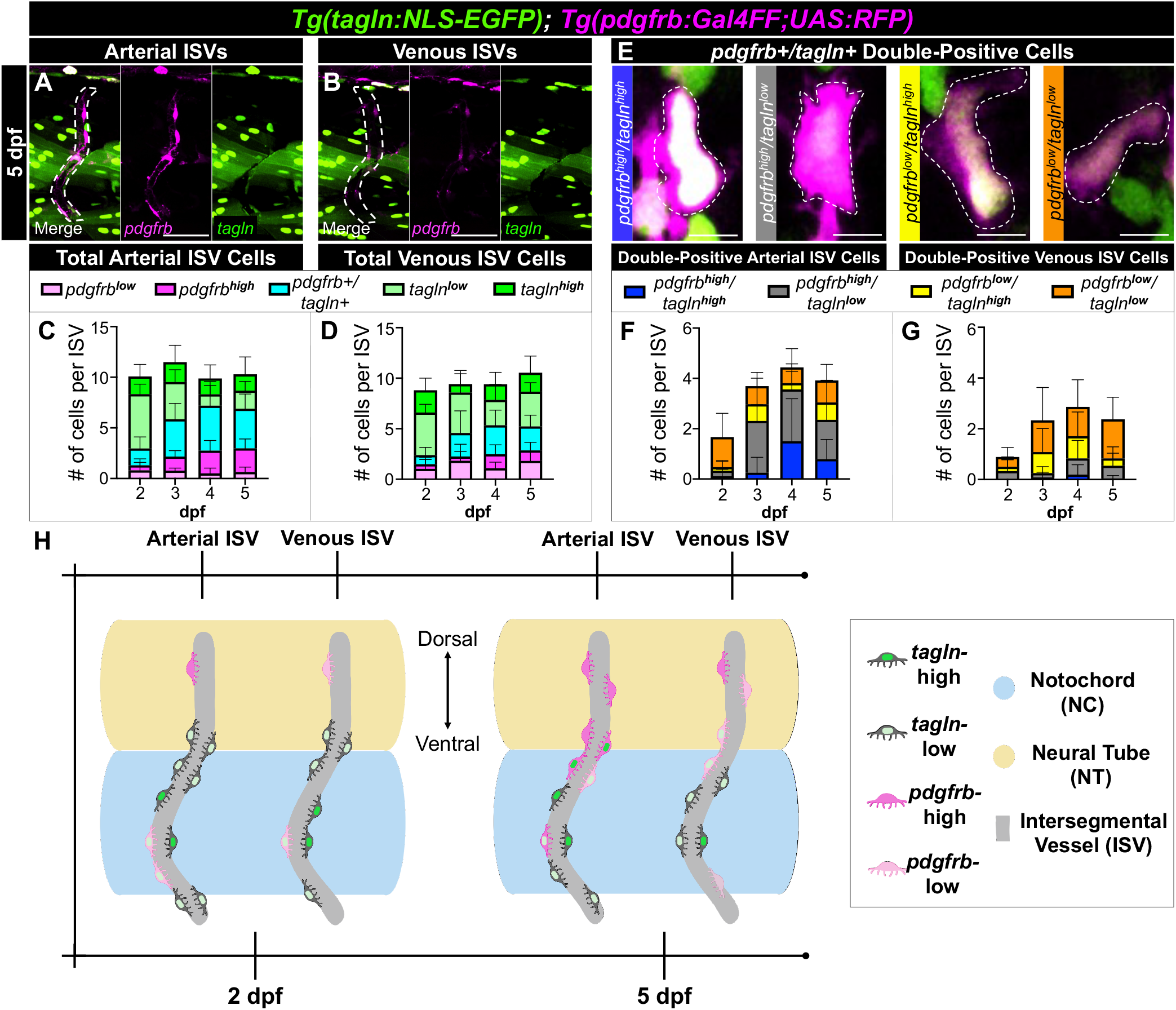
Low and high *pdgfrb*/*tagln* expression is evident along arterial and venous ISVs from 2 to 5 dpf. (A,B) Maximum intensity projection images of arterial and venous ISVs at 5 dpf. ISVs are outlined by white dotted lines. *tagln*-positive skeletal muscle nuclei are particularly evident in the trunk region near the ISVs, thus requiring frame-by-frame analysis to identify *tagln*-positive perivascular cells. (C,D) Bar graph representations of the number of low and high single- and double-positive cells on the arterial (C) and venous (D) ISVs. Arterial ISVs accumulate *pdgfrb*-high mural cells more robustly than venous ISVs. (E) Representative images of the combinations of *pdgfrb*/*tagln* double-positive cells. (F,G) Bar graph representations of the number double-positive cells on the arterial (F) and venous (G) ISVs. *pdgfrb*/*tagln* double-positive cells were more apparent on arterial ISVs. (H) Schematic diagram summarizing the general localization of perivascular cells on the ISVs at 2 and 5 dpf. *pdgfrb* single-positive cells were more likely to be found on the dorsal ISV region, *tagln* single-positive cells primarily near the notochord ISV region, and double-positive cells nearest to the notochord-neural tube interface by 5 dpf. Schematic created with BioRender.com. At least 6 embryos were analyzed at each age. ISVs, intersegmental vessels. Scale bars for A,B: 50 μm; scale bars for E: 5 μm.

From our ISV data, we observed findings similar to that of the perivascular fibroblasts (Rajan et al., 2020): the number of perivascular cells associated with the ISVs remained constant between 2 to 5 dpf, the cells were equally distributed on aISVs and vISVs, and *pdgfrb*-high cells became more apparent on aISVs than vISVs over time (Figs 5A-D; S5A-H). Furthermore, we were able to identify *pdgfrb*/*tagln* double-positive cells with all combinations of high and low expression that increased in proportion up to 4 dpf (Fig. 5E-G). Our data showed that double-positive cells were more common on aISVs than vISVs and, as expected, the double-positive cells on the aISVs were more likely to be *pdgfrb*-high (Fig. 5F,G). *tagln*-high mural cells were constant on all ISVs between 2-5 dpf (Fig. 5C,D), suggesting that on ISVs, mural cells may diverge from the typical expression patterns of differentiation that would predict upregulated *tagln* at late stages in the process of maturation.

*tagln*-low cells were the most common perivascular cell associated with the ISVs at early timepoints (Fig. 5C,D). We speculated that the identity of these cells would overlap with the previously described sclerotome-derived perivascular fibroblast population. Indeed, when using the *twist1a* transgenic line as a sclerotome marker, we identified *twist1a*-positive cells that reside at the notochord-neural tube interface and notochord region as *tagln*-low, even prior to their association with the ISVs (Fig. S6A,D,E). Conversely, a subset of cells that likely originated from the dorsal sclerotome—based on their localization adjacent to the neural tube (Rajan et al., 2020)—were *tagln*-negative (Fig. S6B,C). These data indicate that the ventral sclerotome is likely distinct from dorsal sclerotome by the early and low expression of *tagln*. Additionally, these data reveal that ISV mural cells can express *tagln* as precursors prior to upregulating *pdgfrb*.

We also observed consistent *tagln*/*pdgfrb* expression patterns in mural cells based on their localization on the ISVs. *tagln* single-positive cells were more apparent on the ISVs in the region adjacent to the notochord (Fig. 5H), consistent with previous reports of ventral sclerotome migration to this region (Rajan et al., 2020, Ma et al., 2018). *pdgfrb* single-positive cells were highly associated with the ISVs near to the neural tube (Fig. 5H), where the ISVs are primarily populated by dorsal sclerotome (Rajan et al., 2020). Finally, *pdgfrb*/*tagln* double-positive cells were evident at two sites by 5 dpf—closest to the notochord-neural tube interface (Fig. 5H), which could indicate a site-specific maturation event, and near DA/aISV boundaries, which would suggest that these are vSMCs from the ventral side of the DA that have migrated to the aISVs.

### A unique sclerotome-derived mural cell progenitor population resides along the zebrafish midline

From imaging *tagln*- and *pdgfrb*-positive cells near the ISVs in *Tg(tagln:NLS-EGFP); Tg(pdgfrb:Gal4FF; UAS:RFP)* embryos, we discovered several cell populations with unexpected *pdgfrb*/*tagln* expression near the midline. First, we note the medial floor plate cells are *pdgfrb*/*tagln* double-positive (Fig. 6, MFP cells). Additionally, we observe a *tagln* single-positive population of cells along the midline that accumulates below the medial floor plate cells and above the notochord beginning at 3 dpf (Fig. 6, *tagln*+ midline cells). Finally, we describe a mural cell progenitor population near the notochord-neural tube interface that borders either side of the midline. This population—which begins as *tagln* single-positive cells before acquiring *pdgfrb* expression—can be observed migrating to the nearby ISVs from 2 to 5 dpf (Fig. 6, MC progenitor population).

**Figure 6:**
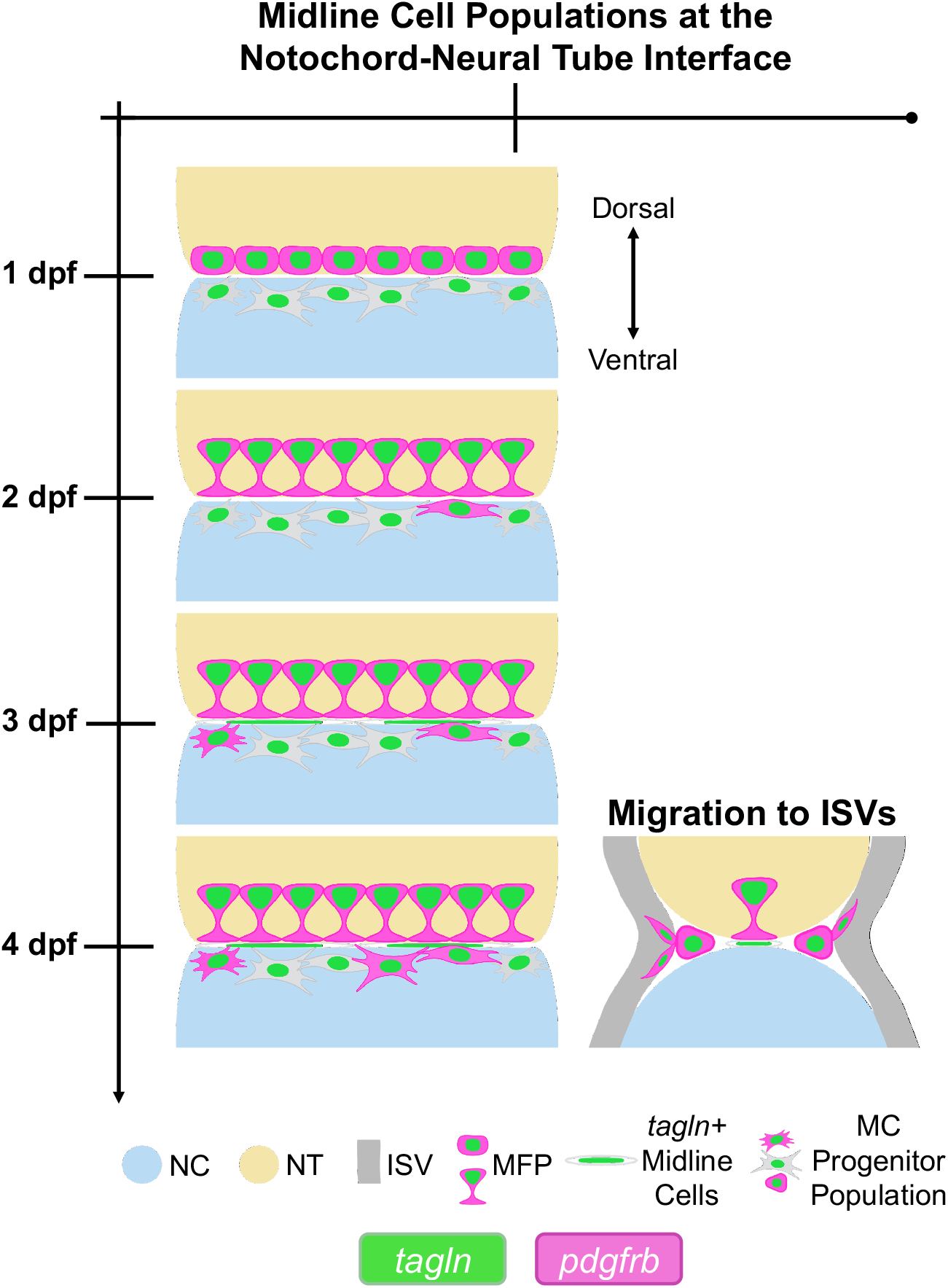
Summary of the midline cell populations at the notochord-neural tube interface. We identify the medial floor plate cells as *pdgfrb*/*tagln* double-positive in early zebrafish development (MFP). Additionally, we observe a *tagln* single-positive population of cells along the midline that accumulates below the medial floor plate cells beginning at 3 dpf (*tagln*+ Midline Cells). Finally, we describe a mural cell progenitor population near the notochord-neural tube interface that borders either side of the midline. This population—which begins as *tagln* single-positive cells before acquiring *pdgfrb* expression—can be observed migrating to the nearby ISVs from 2 to 5 dpf (MC Progenitor Population). Schematic created with BioRender.com. NC, notochord. NT, neural tube. ISV, intersegmental vessel. MFP, medial floor plate. MC, mural cell.

Of the structures that reside at or near the midline, the medial floor plate is one of the most recognizable due to its hourglass-like shape (Zheng et al., 2022). Medial floor plate cells—which reside at the ventral side of the neural tube and contribute to neuronal differentiation (Strähle et al., 2004)—have been previously identified as *pdgfrb*-positive (Ando et al., 2016, Wiens et al., 2010). We observed intense *tagln* co-expression in the medial floor plate cells from 1 to 5 dpf (Fig. 7A-K), indicating that *tagln* may have as-of-yet unidentified developmental functions in primitive, non-muscular tissues.

**Figure 7.**
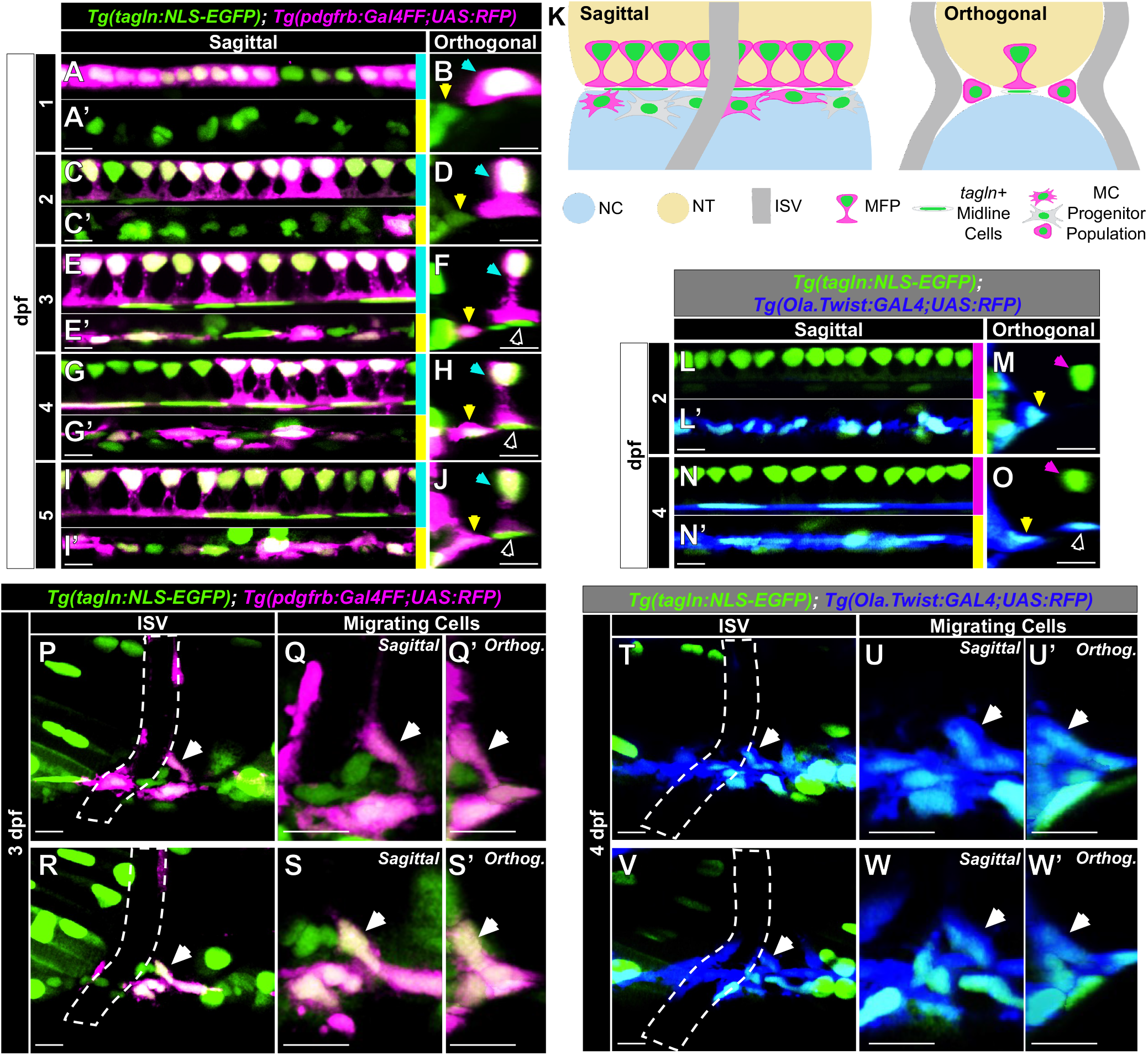
A *tagln*-positive sclerotome-derived mural cell progenitor population at the notochord-neural tube interface upregulates *pdgfrb* and migrates to the ISVs. (A-J) Maximum intensity projection images of cell populations near the notochord-neural tube interface. Sagittal (A,C,E,G,I) and orthogonal (B,D,F,H,J) views of the medial floor plate from 1-5 dpf reveal *pdgfrb*/*tagln* co-expression (cyan arrows). A separate *tagln* single-positive population of cells accumulates immediately below the MFP starting at 3 dpf (E-J, white outlined arrows). Finally, an independent *tagln* single-positive population— hereafter the mural cell progenitor population—aligns near the notochord-neural tube interface (A’-I’,B,D,F,H,J, yellow arrows) and begins to elongate and upregulate *pdgfrb* over time. (K) Schematic diagram illustrating the three described cell populations from a sagittal and orthogonal view. (L-O) The *tagln*-positive midline cells (white outlined arrows) and mural cell progenitors (yellow arrows) are *twist1a*-positive, indicating they are derived from the sclerotome, unlike the medial floor plate (pink arrows). (P-W) Cells from the mural cell progenitor population can be seen migrating to the ISVs using *pdgfrb* and *tagln* as markers (P-S) or *twist1a* and *tagln* (T-W). ISVs are outlined. White arrows indicate migrating cells. Schematic created with BioRender.com. NC, notochord. NT, neural tube. ISV, intersegmental vessel. MFP, medial floor plate. MC, mural cell. Scale bars: 10 μm.

Furthermore, we note a *tagln* single-positive population of elongated cells along the midline beginning at 3 dpf that accumulates between the notochord and neural tube— immediately ventral to the medial floor plate cells (Fig. 7A-K). This population appears to organize as a continuous, thin, rod-like structure that is primarily one cell in width and height. We confirmed that this *tagln*-positive midline cell population originates from the sclerotome—unlike the adjacent medial floor plate—as evidenced by *twist1a* expression (Fig. 7L-O). The function of this previously unidentified structure is currently unknown.

Finally, we observed a cell population—which we will term mural cell progenitors— near the notochord-neural tube interface that borders either side of the midline and migrates to the ISVs (Fig. 7A-W; Movies 2,3). At 1 dpf, these cells expressed *tagln*, but not *pdgfrb* (Fig. 7A’), and at 2 dpf, a fraction of these cells began to express *pdgfrb* (Fig. 7C’). Over the next 3 days these cells elongated and increased *pdgfrb* expression until approximately half were *pdgfrb*/*tagln* double-positive at 5 dpf (Fig. 7E’,G’,I’).

Due to this mural cell progenitor population’s proximity to the notochord and floor plate, we questioned whether they originated from or resided within either of these structures. Using a sonic hedgehog signaling molecule a (*shha*) transgenic line to label the floor plate (Shkumatava et al., 2004, Ertzer et al., 2007), we were able to visualize the mural cell progenitor population as sitting outside of the neural tube (Fig. S7A,B). Next, we used a procollagen, type IX, alpha 2 (*col9a2*) transgenic line to visualize notochord sheath cells in relation to the progenitor population (Garcia et al., 2017). Notochord sheath cells are reported to express *tagln* in the notochord bud during regeneration (Sinclair et al., 2021); therefore, we wanted to determine if the sheath cells may be the source of the mural cell progenitor population. However, the progenitor population did not overlap with notochord sheath cells (Fig. S7C,D), though we confirm that sheath cells do express low levels of *tagln* during development (Fig. S7E). Jointly, these data showed that the mural cell progenitor population abuts the neural tube and notochord, but does not overlap with these structures.

We next used the sclerotome-marker *twist1a* and the neural crest-marker SRY-box transcription factor 10 (*sox10*) transgenic lines (Lee et al., 2013, Kucenas et al., 2008) to test the origins of this novel mural cell progenitor population. We identified this progenitor population as exclusively derived from *twist1a*-positive sclerotome and not *sox10*-positive neural crest (Figs 7L-O; S7F,G; Movie 3). Since Notch signaling from arterial endothelium has been shown to be involved in maturation of mural cell progenitors to *pdgfrb*-high mural cells (Ando et al., 2019), we speculated that the progenitor population could be in contact with a blood vessel along the midline that would induce an increase in *pdgfrb* expression. However, there is no blood vessel present along the midline at this time point (Fig. S7H), nor do cells that increase *pdgfrb* expression in the progenitor population reside near blood vessels (Fig. S7I,J). This suggests that the induction of *pdgfrb* expression in this mural cell progenitor population is not controlled by contact with the arterial vasculature.

Using *tagln*, *pdgfrb*, and *twist1a* as cellular markers, we observed cells from this progenitor population migrate to the ISVs along the region adjacent to the notochord-neural tube interface (Fig. 7P-W; Movies 2,3). Progenitor cells migrated to both aISVs and vISVs from 2 to 5 dpf. Due to the location of this mural cell progenitor population in relation to the ISVs, a proportion of the *pdgfrb*/*tagln* double-positive cells that we observe on ISVs in the region closest to the notochord-neural tube interface (Fig. 5H) likely originate from this progenitor source.

## Discussion

### *pdgfrb* and *tagln* transgenic zebrafish reporters

In this study, we use *Tg(pdgfrb:Gal4FF; UAS:RFP)* and *Tg(tagln:NLS-EGFP)* transgenic lines to identify single- and double-positive pericytes/vSMCs in the cranial, axial, and ISVs at 1 to 5 dpf. Our nuclear-GFP *tagln* reporter facilitated identification of novel *tagln*-positive cells in the zebrafish trunk. High *tagln* expression is evident in skeletal muscle, which has made it difficult in the past to explore *tagln* expression patterns in cells near to the skeletal muscle with cytosolic-GFP *tagln* reporters. With the nuclear-GFP *tagln* reporter, we have revealed *tagln* expression in the hypochord, the medial floor plate, and in sclerotome-derived mesenchyme in the trunk region.

To rule out whether these expression patterns were an artifact of the nuclear-GFP *tagln* reporter, we sought to confirm *tagln* expression in these structures using an independently generated cytosolic-GFP *tagln* reporter (*Tg(tagln:EGFP)*; Fig. S8A-H). As expected, GFP expression in the skeletal muscle decreased the efficiency and resolution of identifying *tagln*-positive cells in the trunk, though we were able to visualize GFP expression in the hypochord, ISV mural cells, the novel mural cell progenitor population we describe, the medial floor plate, and *tagln*-positive midline cells. Thus, despite high background GFP signal, we validate *tagln* expression in these structures using the cytosolic-GFP *tagln* reporter.

The *pdgfrb* reporter line—which utilizes the Gal4/UAS system (Ando et al., 2016)— presented with occasional RFP expression mosaicism. This mosaicism, when present, was most apparent in the medial floor plate cells (Fig. 7A-J), though it is difficult to assess the penetrance at which other structures were affected. Due to this, there may be *pdgfrb*-positive cells that are underrepresented in this study, particularly cells that express low levels of *pdgfrb*. However, despite potential mosaicism issues, the expression patterns that we visualized over time and in various regions were consistent across all samples.

### Cranial Mural Cells

Previous reports show that *pdgfrb*-positive pericytes are the predominant mural cell in the mid- and hindbrain prior to 5 dpf, at which point arterial-associated mural cells begin to express *tagln* and other vSMC markers (Ando et al., 2019, Ando et al., 2016, Ando et al., 2021a, Whitesell et al., 2014, Wang et al., 2014). We observed similar findings, in that *pdgfrb* single-positive pericytes were the most prevalent type of mural cell along embryonic cranial vessels, with eventual *tagln* co-expression evident—yet sparsely represented—in a subset of *pdgfrb*-positive arterial mural cells at 4 dpf. No *tagln* single-positive cells were detected at the time points assessed. Based on these data, and supported by the findings of previous studies, we conclude that *pdgfrb* is an ideal reporter to visualize cranial mural cell association with blood vessels up to 5 dpf, with the caveat that a large proportion of early *pdgfrb*-positive cranial cells are neural crest or mesenchyme progenitors that are not associated with vessels.

### Mural Cells of the Axial Vessels

In the axial vessels (i.e. DA, PCV, CA, and CV), we corroborate the previous findings regarding vSMC expression patterns on the ventral side of the DA (Ando et al., 2021a, Stratman et al., 2017, Ando et al., 2016, Ando et al., 2019). Specifically, we detect *pdgfrb* single-positive cells at 2 dpf, where they then expand in number and increase expression of *tagln* starting at 3 dpf (Fig. 2). Moreover, we observe a slow and sporadic increase of *pdgfrb* single-positive cells on the PCV between 1 to 5 dpf (Fig. 2). At 5 dpf, there are twice as many *pdgfrb*-positive mural cells associated with the ventral side of the DA than the entire PCV, which highlights the clear difference in the magnitude of mural cell recruitment between arteries and veins early in development.

Our analysis of both the anterior and caudal axial vessels also reveals that accumulation and maturation of these mural cells occurs in an anterior to posterior fashion. By 1 dpf, the anterior and posterior regions of the main axial vessels have formed, somite segmentation has been completed, and the sclerotome is distinct from other somitic tissues (Stickney et al., 2000). Despite this, DA vSMCs—which are mainly derived from the sclerotome (Stratman et al., 2017)—mature one to two days earlier than the vSMCs of the CA (Fig. 2). There may be several reasons for this delay, with one hypothesis being that the DA provides earlier signals than the CA to initiate maturation of vSMCs. Alternatively, since somites develop in an anterior to posterior wave (Stickney et al., 2000), the age of the somites from which mural cells are derived could influence the timing of vSMC maturation, even after vSMCs have accumulated on the vessels. More work will be needed in this area to resolve the cues driving this temporal differentiation pattern.

### The Hypochord as a Source of *tagln*-Positive Cells

Through use of the nuclear-GFP *tagln* reporter, we found that the hypochord contributes to *tagln*-positive cells along the dorsal side of the DA up to at least 5 dpf (Fig. 2). Interestingly, the hypochord and medial floor plate—which we also show is *tagln*-positive—are among the earliest structures in the zebrafish embryo to co-express *pdgfrb* and *tagln*. The hypochord, floor plate, and notochord have been shown to be derived from the multipotent midline progenitor cells in the dorsal organizer or the tailbud (Row et al., 2016, Latimer and Appel, 2006)—a common origin which may, in part, explain how two structures that are dissimilar in function share this unique expression pattern. Furthermore, in the invertebrate *Ciona*, researchers show that a series of calponin/transgelin family genes (*tagln-r.b, tagln-r.c, tagln-r.d, tagln-r.e*) are expressed in the *Ciona* floor plate, notochord, and endodermal strand (the *Ciona* analogue for the hypochord), indicating that Tagln expression in midline structures may have a conserved role in embryogenesis (Oonuma et al., 2021). As Tagln is known primarily as an actin-binding protein in contractile cells, the role of Tagln in these non-contractile, non-muscular, primitive midline structures remains unknown.

The persistence of the elongated *tagln*-positive cells on the dorsal side of the DA indicates that the cells of the hypochord are not entirely transient, as previously assumed. While previous studies have observed hypochord cells undergoing apoptosis (Löfberg and Collazo, 1997, Cleaver et al., 2000), we propose that a subset of hypochord cells survive and reprogram their cellular function and morphology after the DA has been appropriately oriented. Due to the maintained expression of *tagln* and proximity to the DA, we propose these cells may undertake a mural cell role; however, further research is needed to determine whether these cells function similarly to other vSMC populations.

As the hypochord is present exclusively in fish and amphibians, we questioned whether there is an analogous mammalian structure that could contribute to Tagln-positive cells on the DA of amniotes. Previous reports speculate that the dorsal endoderm—a midline structure that originates from the midline precursor cell population and forms ventral to the notochord (Peyrot et al., 2011)—may play a role in angioblast and DA formation (Poole et al., 2001), similar to the hypochord. However, while the endoderm gives rise to the intestines, lung, liver, and other organs (Wlizla and Zorn, 2015), there are no known reports of endoderm contributing to mural cell populations in amniotes (Holm et al., 2018).

In the mouse embryo, the dorsal and ventral sides of the DA are populated by vSMCs from different sources: the ventral side is first populated by cells from the lateral plate mesoderm, with dorsal vSMCs originating from the somites. Over time, the vSMCs from the lateral plate mesoderm disappear, and the ventral side then becomes populated with somitic vSMCs (Wasteson et al., 2008). While this process is not identical on the zebrafish DA—given that somitic vSMCs are the initial populators of the ventral side of the DA in zebrafish—it does indicate that dorsal and ventral vSMCs can arise from divergent origins and supports that this could be a conserved process.

### Mural Cells of the Intersegmental Vessels

In the ISV region, we note perivascular cells associated with the vessels exhibiting all combinations of high and low expression of *pdgfrb* and *tagln*. To summarize, *pdgfrb*-high cells were more common on arterial than venous ISVs and increased in number over time. *pdgfrb*/*tagln* double-positive cells were also more common on aISVs and were more likely to be *pdgfrb*-high on aISVs than vISVs. *tagln*-high cells were constant in number between 2 to 5 dpf, and *tagln*-low cells—the most common expression pattern represented—were revealed to be sclerotome-derived cells that were migrating to or associated with the ISVs. Further, expression patterns were related to mural cell location on the ISVs, with *pdgfrb* single-positive cells more likely to be found on the dorsal ISV region, double-positive cells nearest to the notochord-neural tube interface, and *tagln* single-positive cells primarily near the notochord ISV region by 5 dpf (Fig. 5H).

These data indicate that *tagln* may not exclusively label mature mural cells, but instead could also label a subset of mural cell precursors in the trunk. These precursors likely overlap with the perivascular fibroblast population described by Rajan et al (Rajan et al., 2020), as they are both sclerotome-derived and populate the ISVs prior to reported mural cell maturation. This would suggest that a subset, but not all, of the *tagln*-positive early precursors on the ISVs will mature into mural cells.

### Sclerotome-Derived *tagln*-Positive Cell Populations in the Trunk

Use of our nuclear-GFP *tagln* reporter allowed us to ascertain that sclerotome-derived cells in the zebrafish trunk are primarily *tagln*-positive as early as 1 dpf. Due to the main localization of these *tagln*-positive cells at the notochord-neural tube interface or below, we speculate that the *twist1a*/*tagln* double-positive cells arise from the ventral sclerotome, with *twist1a*-positive/*tagln*-negative cells descending from the dorsal sclerotome. Of note, there are two unique sites where sclerotome-derived *tagln*-positive cells accumulate during development.

First, we identified a *twist1a*/*tagln* double-positive midline population between the neural tube and notochord. This population begins to accumulate beneath the medial floor plate cells at 3 dpf and exhibits high *tagln* expression (Fig. 6, *tagln*+ midline cells). Though this structure’s function is unknown, its location would suggest that it may exist as a cushion or as structural support to the notochord-neural tube interface.

Second, we identified a mural cell progenitor population that resides adjacent to the midline at the notochord-neural tube interface. This *tagln*-positive population upregulates *pdgfrb* expression sporadically in an increasing number of cells between 2 to 5 dpf and migrates to the ISVs (Fig. 6, MC progenitor population; Movies 2,3). Due to the location of this mural cell progenitor population in relation to the ISVs, we suspect that a proportion of the *pdgfrb*/*tagln* double-positive cells that we observe on ISVs in the region closest to the notochord-neural tube interface likely originate from this progenitor source. Uniquely, the upregulation of *pdgfrb* expression in this mural cell progenitor population occurs independently of contact with arterial endothelium. We hypothesize that upregulation of *pdgfrb* confers motility cues to the mural cell progenitors to then migrate to blood vessels; however, further studies are needed to determine the mechanisms behind this seemingly stochastic *pdgfrb* upregulation.

This progenitor population also demonstrates that mural cell precursors can express *tagln* prior to upregulating *pdgfrb*. Since *pdgfrb* is associated with precursor populations and *tagln* is regarded as a mature vSMC marker, this finding suggests that the expression patterns related to mural cell maturation are more nuanced than previously reported. Therefore, we ask: what role could Tagln play prior to Pdgfrb upregulation? Several studies report that Tagln assists in cell motility: loss of Tagln decreases motility (Elsafadi et al., 2016), whereas overexpression leads to pathological motility in situations such as metastatic cancer (Zhou et al., 2016, Chen et al., 2019). Since sclerotome cells are inherently migratory as they populate and form various tissues, this could help explain the early expression of Tagln in mural cell precursors during embryogenesis. Future work in this area will continue to reveal exciting new findings driving site-specific and temporal regulation of mural cell biology.

## Materials and Methods

### Zebrafish

Zebrafish (*Danio rerio*) embryos were raised and maintained as described (Westerfield, 2000, Kimmel et al., 1995). Zebrafish husbandry and research protocols were reviewed and approved by the WUSTL Animal Care and Use Committee. Established transgenic lines that were used in this study are listed in the table below.

| Genotype | Allele | Origin |
| --- | --- | --- |
| *Tg(-2.4shha-ABC:GFP)* | *sb15* | (Shkumatava et al., 2004) |
| *Tg(col9a2:mCherry)* | *pd1150* | (Garcia et al., 2017) |
| *Tg(kdrl:mCherry-CAAX)* | *y171* | (Fujita et al., 2011) |
| *Tg(Ola.Twist:GAL4)* | *sq11* | (Lee et al., 2013) |
| *Tg(sox10:mRFP)* | *vu234* | (Kucenas et al., 2008) |
| *Tg(tagln:eGFP)* | *p151* | (Seiler et al., 2010) |
| *Tg(tagln:NLS-EGFP-2A-CFP-FTASE)* | *y450* | (Stratman et al., 2017) |
| *Tg(5xUAS:RFP)* | *zf83* | (Asakawa et al., 2008) |
| *TgBAC(pdgfrb:Gal4FF)* | *ncv24* | (Ando et al., 2016) |

### Image Acquisition and Processing

Embryos and larvae were anesthetized in buffered 0.2 mg/mL Tricaine-S (Western Chemical, Inc; NC0872873) and mounted in 0.8% low-melting point agarose (IBI Scientific; #IB70056) on a 35-mm glass-bottom petri dish (MatTek; #P35G-1.5-20-C). Confocal images were obtained using a 4x or 40x objective with a W1 Spinning Disk confocal microscope, a Fusion camera, and the Nikon Eclipse Ti2-E base. Images were then deconvolved using Nikon’s NIS-Elements software. Fiji image processing software was used for counting number of cells per vessel length and for generating final images. Embryos were consistently imaged in the afternoon hours, thus: ‘1 dpf’ represents embryos between 28-34 hours post-fertilization (hpf); ‘2 dpf’ represents 52-58 hpf; ‘3 dpf’ represents 76-82 hpf; ‘4 dpf’ represents 100-106 hpf; ‘5 dpf’ represents 124-130 hpf. At least 6 embryos were imaged and analyzed per developmental timepoint.

## Acknowledgements

We thank Tony Tsai’s lab for the *Tg(−2.4shha-ABC:GFP)* line, Shawn Burgess’s lab for the *Tg(col9a2:mCherry)* line, and Mayssa Mokalled’s lab for the *Tg(sox10:mRFP)* line. Figure schematics were generated using BioRender.com.

## Figure Legends

**Supplementary Figure 1.**
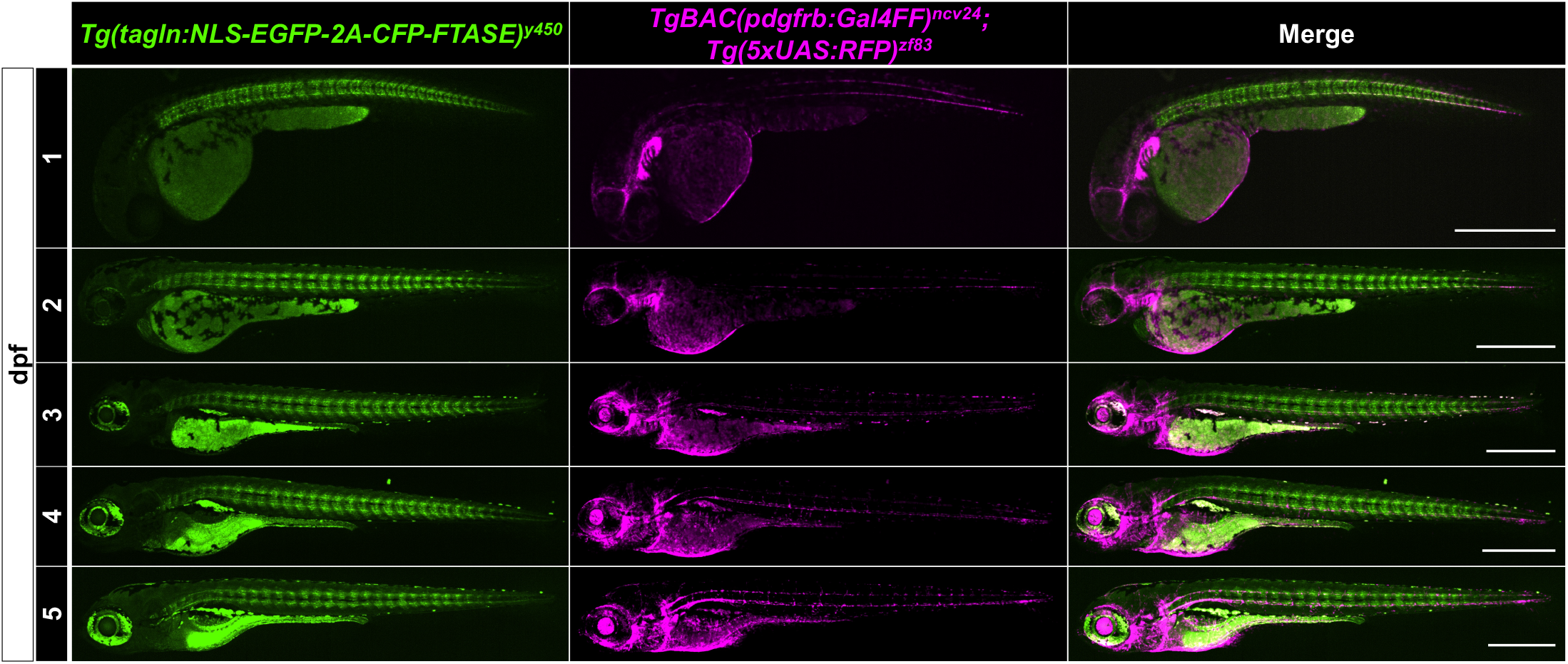
*pdgfrb* and *tagln* fluorescent reporter lines. Lateral view confocal images of *Tg(tagln:NLS-EGFP-2A-CFP-FTASE); TgBAC(pdgfrb:Gal4FF); Tg(5xUAS:RFP)* triple transgenic zebrafish embryos from 1-5 dpf. *tagln* expression becomes increasingly evident in the skeletal muscle of the trunk and the intestinal compartment. *pdgfrb* expression is primarily visible in the anterior and cranial regions. Scale bars: 500 µm.

**Supplementary Figure 2.**
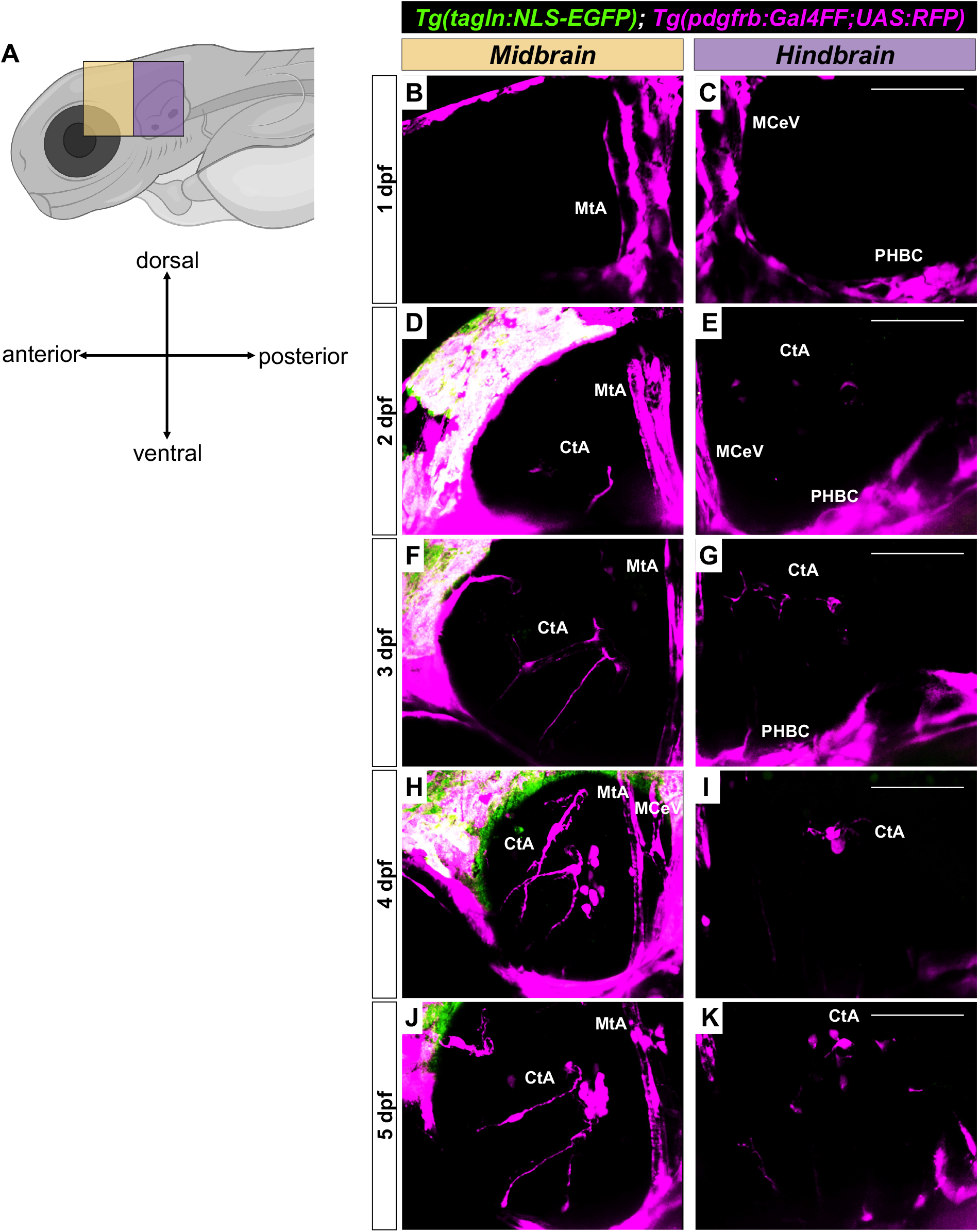
Sagittal views of the mid- and hindbrain of 1 through 5 dpf *Tg(tagln:NLS-EGFP); Tg(pdgfrb:Gal4FF; UAS:RFP)* zebrafish. (A) Schematic diagram illustrating the orientation for imaging sagittal sections of the midbrain (yellow) and hindbrain (purple). (B-K) Maximum intensity projection images of the mid- and hindbrain from 1 to 5 dpf. At 1 dpf (B,C) *pdgfrb*-positive cells are associated with the PHBC, MtA, and MCeV. At 2 dpf (D,E) *pdgfrb*-positive cells begin to accumulate on the CtAs. *pdgfrb*-positive cells continue to accumulate on the CtAs from 3 to 5 dpf (F-K) in both the mid- and hindbrain regions. *tagln*-positive cells were not evident in any of the sagittal images of the mid- or hindbrain. Schematic created with BioRender.com. PHBC, primordial hindbrain channel. MtA, metencephalic artery. MCeV, middle cerebral vein. CtA, central artery. Scale bars: 50 µm.

**Supplementary Figure 3.**
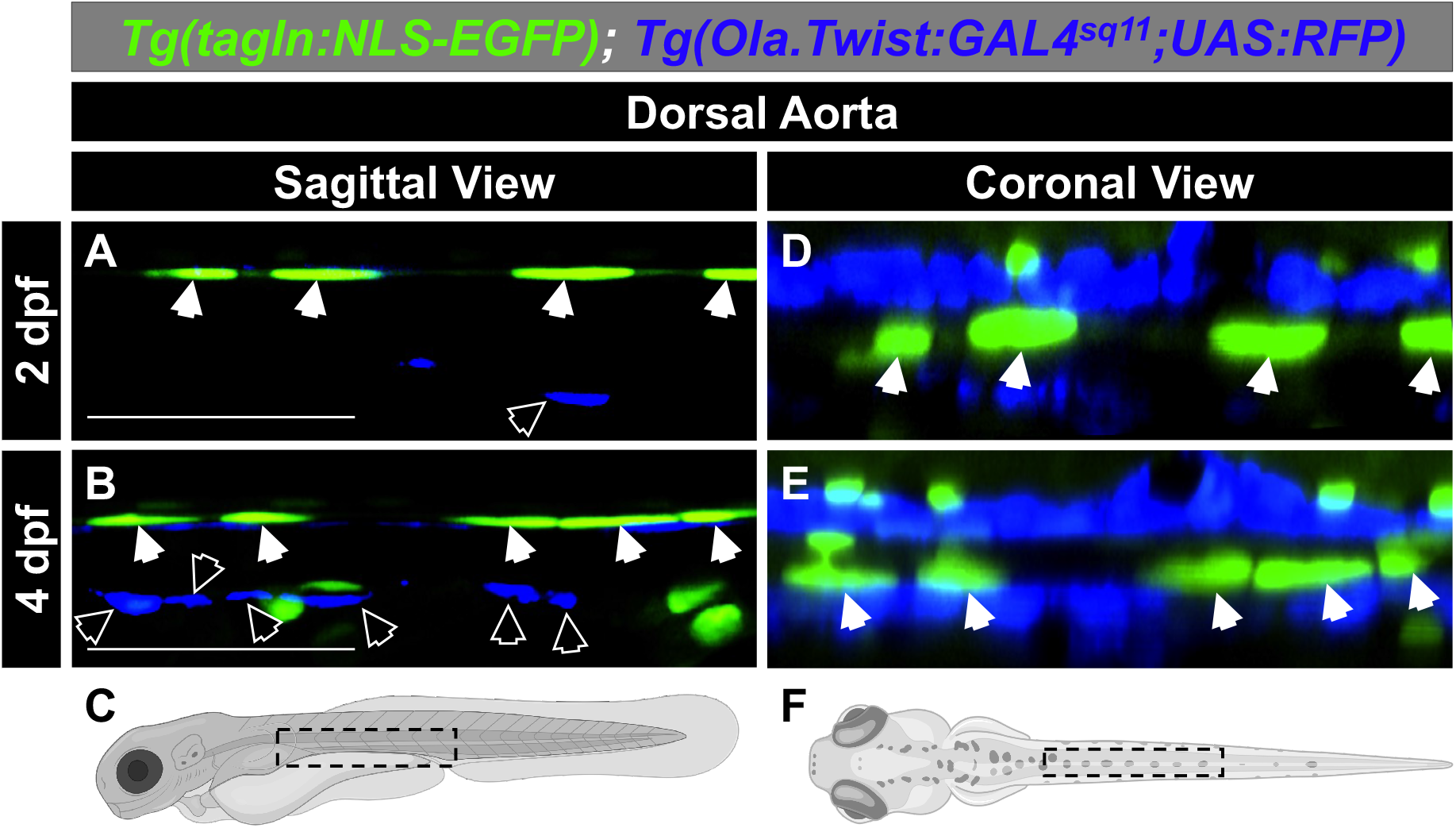
*tagln* single-positive cells on the dorsal side of the DA are not sclerotome-derived. Confocal images of the DA in *Tg(tagln:NLS-EGFP); Tg(Ola.Twist:GAL4; UAS:RFP)* fish at 2 dpf (A,D) and 4 dpf (B,E). From the sagittal view (A-C), *tagln*-positive nuclei derived from the hypochord are apparent at 2 dpf and do not express *twist1a* at the 2 or 4 dpf timepoints (solid white arrows), unlike vSMCs on the ventral side of the DA that do express *twist1a* (white outlined arrows). From the coronal view (D-F), *tagln* single-positive nuclei of the hypochord on the dorsal side of the DA are *twist1a*-negative at both time points (solid white arrows), though *twist1a*-positive cells do abut the DA in this region. These data indicate that the *tagln*-single positive hypochord cells on the dorsal side of the DA are not replaced by sclerotome-derived vSMCs after 2 dpf when the hypochord has been previously reported to disappear. (C,F) Schematic diagrams illustrating the orientation for imaging sagittal (C) or coronal (F) sections of the DA. Schematics created with BioRender.com. DA, Dorsal Aorta. Scale bars: 50 µm.

**Supplementary Figure 4.**
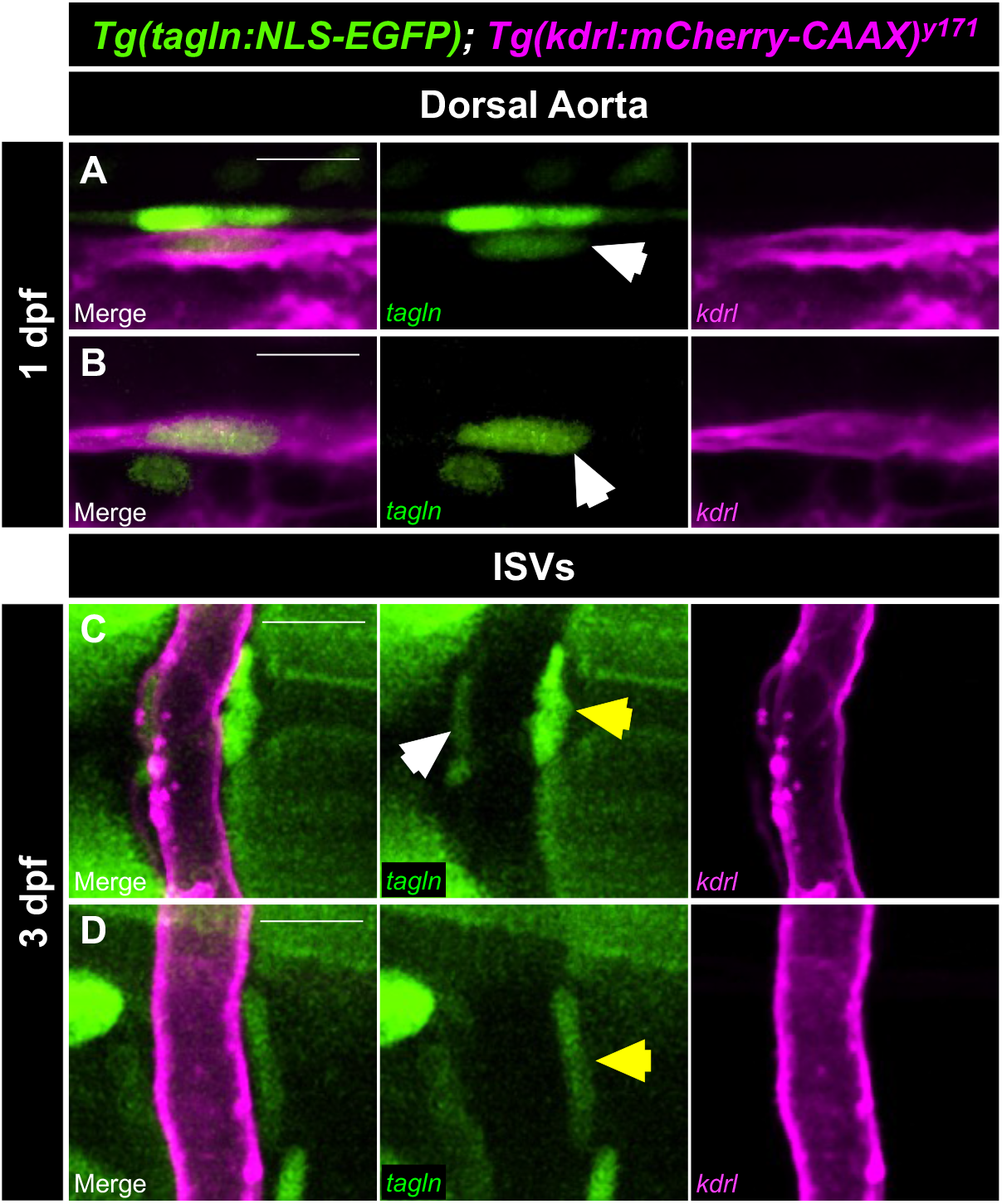
Low *tagln* expression is evident in endothelial cells and a subset of perivascular cells on the ISVs. (A-D) Confocal images of *Tg(tagln:NLS-EGFP); Tg(kdrl:mCherry-CAAX)* fish at 1 dpf (A,B) and 3 dpf (C,D) where *kdrl* is used to mark endothelial cells (magenta). (A-C) Low *tagln* expression is seen in endothelial cells of the aorta and ISVs (white arrows). High (C) and low (D) *tagln* expression is also visible in perivascular cells on the ISVs (yellow arrows). Low *tagln* expression in the endothelial cells of the axial vessels is easily distinguishable from high expression in vSMCs, therefore it did not confound the mural cell counts of the axial vessels of *Tg(tagln:NLS-EGFP); Tg(pdgfrb:Gal4FF; UAS:RFP)* fish. However, ISV endothelial cells and *tagln*-low perivascular cells were difficult to distinguish, thus a small number of endothelial cells may be included in the count of *tagln*-low cells of the ISVs. ISVs, intersegmental vessels. Scale bars: 10 µm.

**Supplementary Figure 5.**
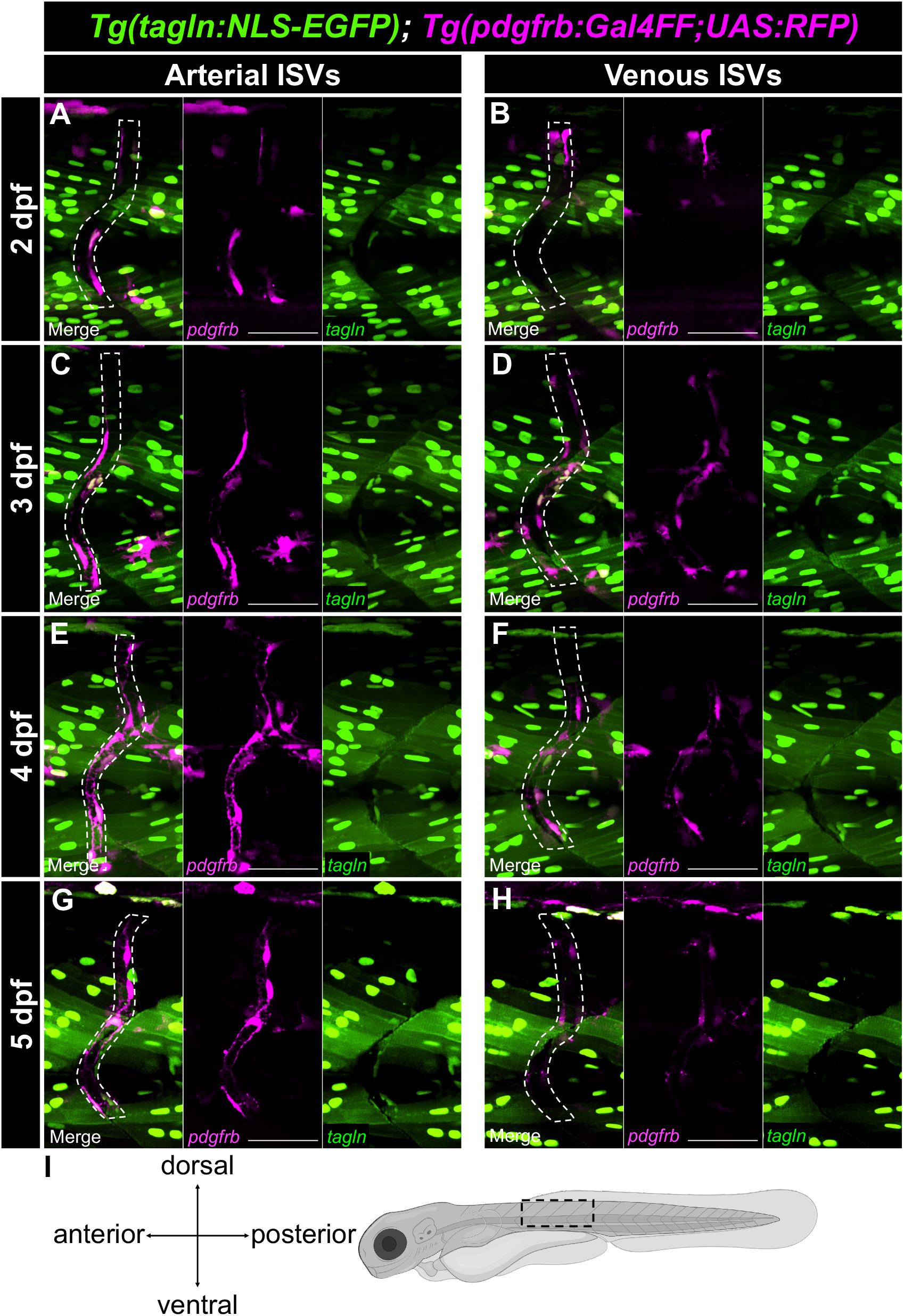
*tagln* and *pdgfrb* expression along arterial and venous ISVs of 2 to 5 dpf *Tg(tagln:NLS-EGFP); Tg(pdgfrb:Gal4FF; UAS:RFP)* zebrafish. (A-H) Maximum intensity projection images of arterial and venous ISVs from 2 to 5 dpf. ISVs are outlined by white dotted lines. *tagln*-positive skeletal muscle nuclei are particularly evident in the trunk region near the ISVs, thus requiring frame-by-frame analysis to identify *tagln*-positive perivascular cells. Arterial ISVs (A,C,E,G) accumulate *pdgfrb*-high mural cells at higher levels than venous ISVs (B,D,F,H). All combinations of low and high *pdgfrb* and *tagln* expression were detected in single- and double-positive perivascular cells surrounding the ISVs at all timepoints, though *pdgfrb/tagln* double-positive cells were more apparent on arterial ISVs. (I) Schematic diagram illustrating the imaged region. Schematic created with BioRender.com. ISVs, intersegmental vessels. Scale bars: 50 µm.

**Supplementary Figure 6.**
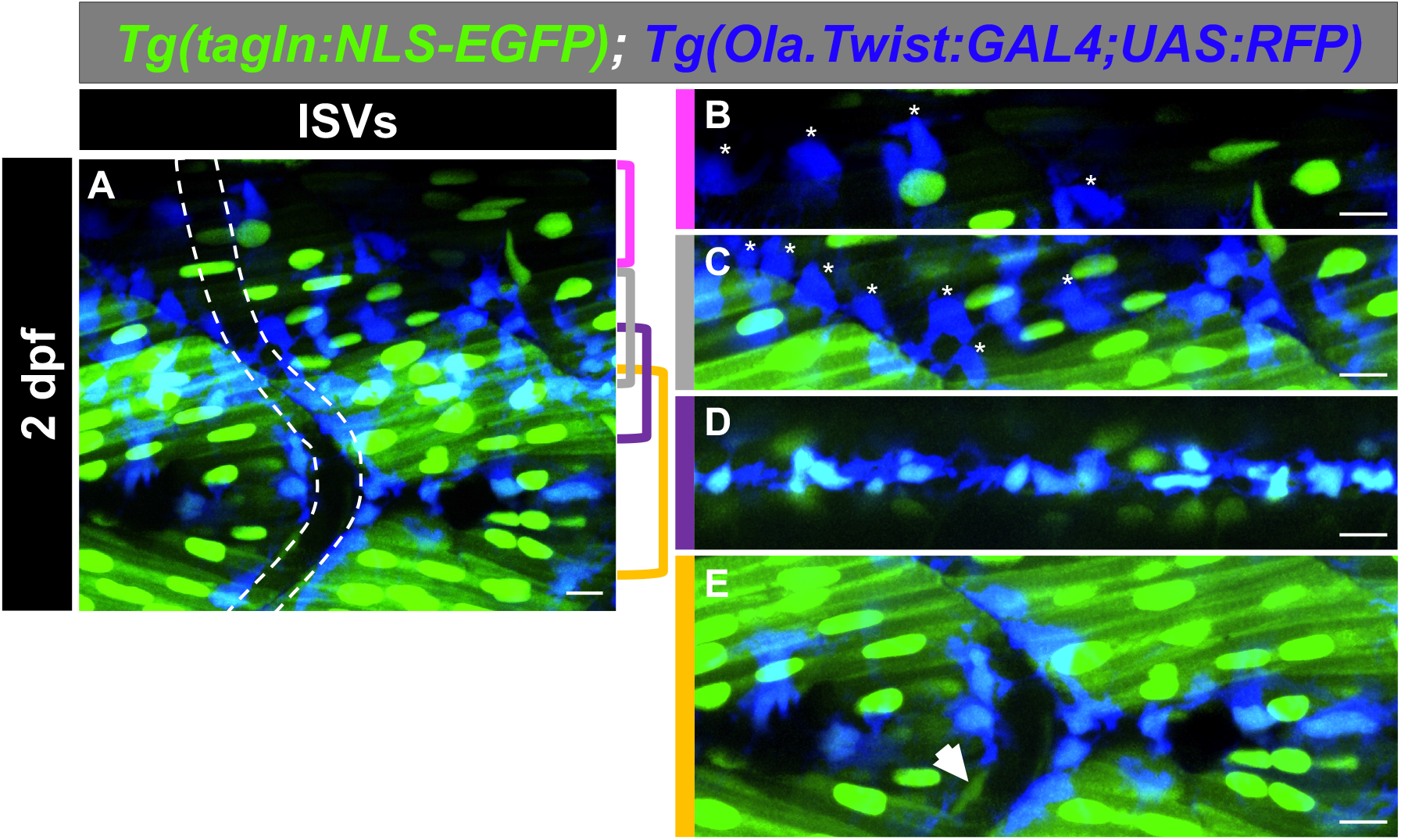
Sclerotome-derived cells with low *tagln* expression are predicted to arise from the ventral sclerotome compartment. (A) Maximum intensity projection image of the ISV region of a *Tg(tagln:NLS-EGFP); Tg(Ola.Twist:GAL4; UAS:RFP)* fish at 2 dpf. ISV is outlined with a white dotted line. (B-E) Discrete regions of panel A are max projected for better visualization of *twist1a*-positive cells near the neural tube (B,C), notochord-neural tube interface (D), and notochord (E). The majority of *twist1a*-positive cells near the neural tube are *tagln*-negative (indicated by stars), while all *twist1a*-positive cells at the notochord-neural tube interface and below are *tagln*-positive. Due to the ventral localization of *tagln/twist1a* double-positive cells, we suspect that these cells arise from the ventral sclerotome, with *twist1a*-single positive cells near the neural tube arising from the dorsal sclerotome. The *tagln*-positive/*twist1a*-negative vascular cell in panel E is likely endothelium (white arrow). ISVs, intersegmental vessels. Scale bars: 10 µm.

**Supplementary Figure 7.**
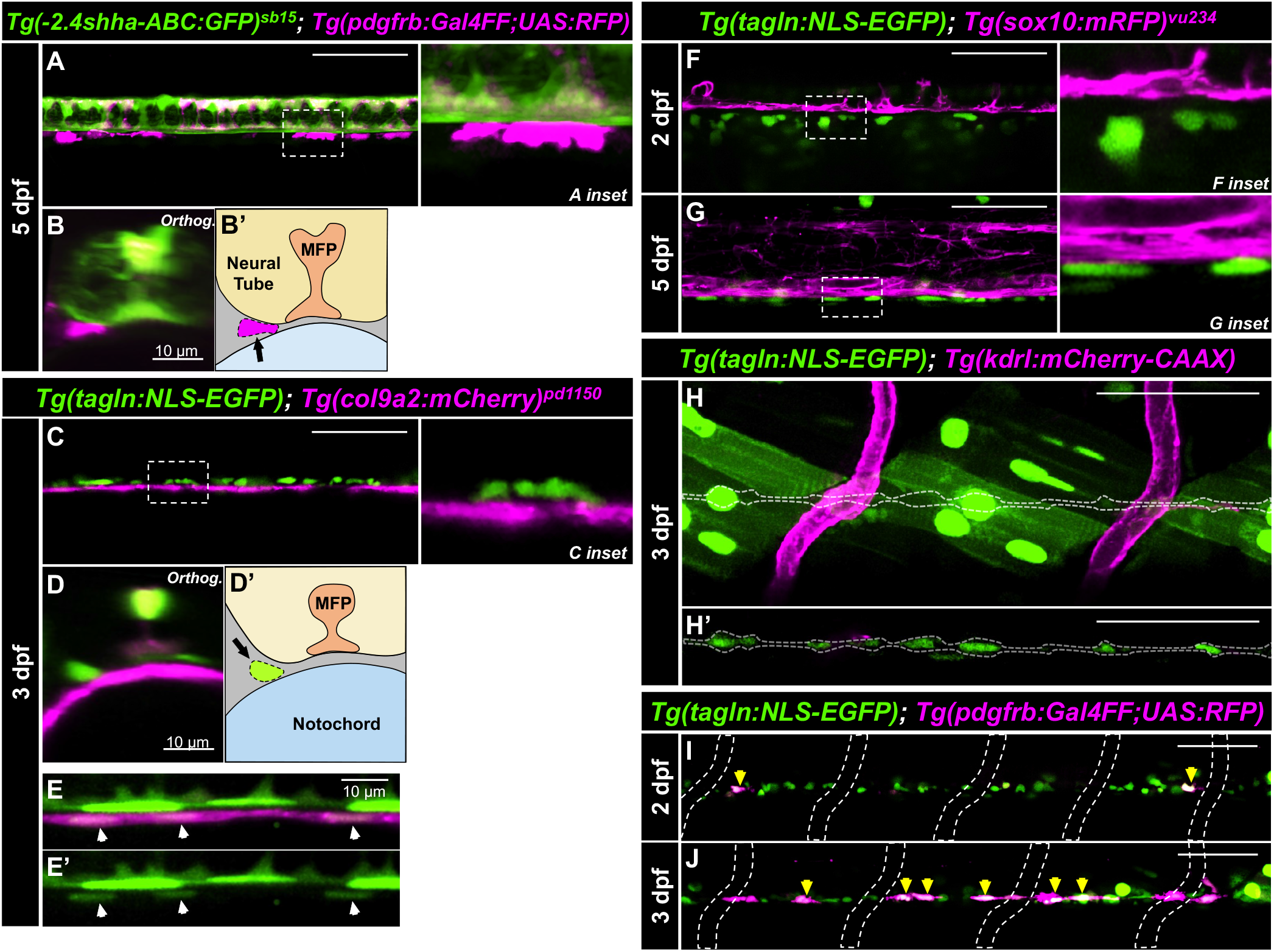
The MC progenitor population resides between the notochord and neural tube, is not derived from neural crest, and does not need to contact endothelium to upregulate *pdgfrb*. (A,B) Confocal images of *Tg(−2.4shha-ABC:GFP); Tg(pdgfrb:Gal4FF; UAS:RFP)* zebrafish to visualize the medial floor plate cells (*shha*) and the MC progenitor population (*pdgfrb*) at 5 dpf. The sagittal (A) and orthogonal (B) views show the MC progenitors outside of the neural tube. (B’) Schematic detailing the structures in panel B. (C-E) Confocal images of *Tg(tagln:NLS-EGFP); Tg(col9a2:mCherry)* zebrafish to visualize the notochord sheath cells (*col9a2*) and the MC progenitor population (*tagln*) at 3 dpf. The sagittal (C) and orthogonal (D) views show the MC progenitors outside of the notochord. (D’) Schematic detailing the structures in panel D. (E,E’) Notochord sheath cells (white arrows) express low levels of *tagln*. (F,G) Confocal images of *Tg(tagln:NLS-EGFP); Tg(sox10:mRFP)* fish to visualize the neural crest (*sox10*) and the MC progenitor population (*tagln*) at 2 dpf (F) and 5 dpf (G). MC progenitor cells do not overlap with neural crest populations. (H) Confocal images of *Tg(tagln:NLS-EGFP); Tg(kdrl:mCherry-CAAX)* fish to visualize endothelial cells (*kdrl*) and the MC progenitor population (*tagln*) at 3 dpf. The MC progenitor population (outlined with a white dotted line) is not adjacent to a blood vessel. (H’) Z-stack slice showing the MC progenitor population from panel H without the skeletal muscle. (I,J) Confocal images of *Tg(tagln:NLS-EGFP); Tg(pdgfrb:Gal4FF; UAS:RFP)* fish at 2 dpf (I) and 3 dpf (J). ISVs are outlined with white dotted lines. MC progenitor cells upregulate *pdgfrb* expression (yellow arrows) without contact with the endothelium. MC, mural cell. MFP, medial floor plate. Scale bars: 50 µm unless otherwise indicated.

**Supplementary Figure 8.**
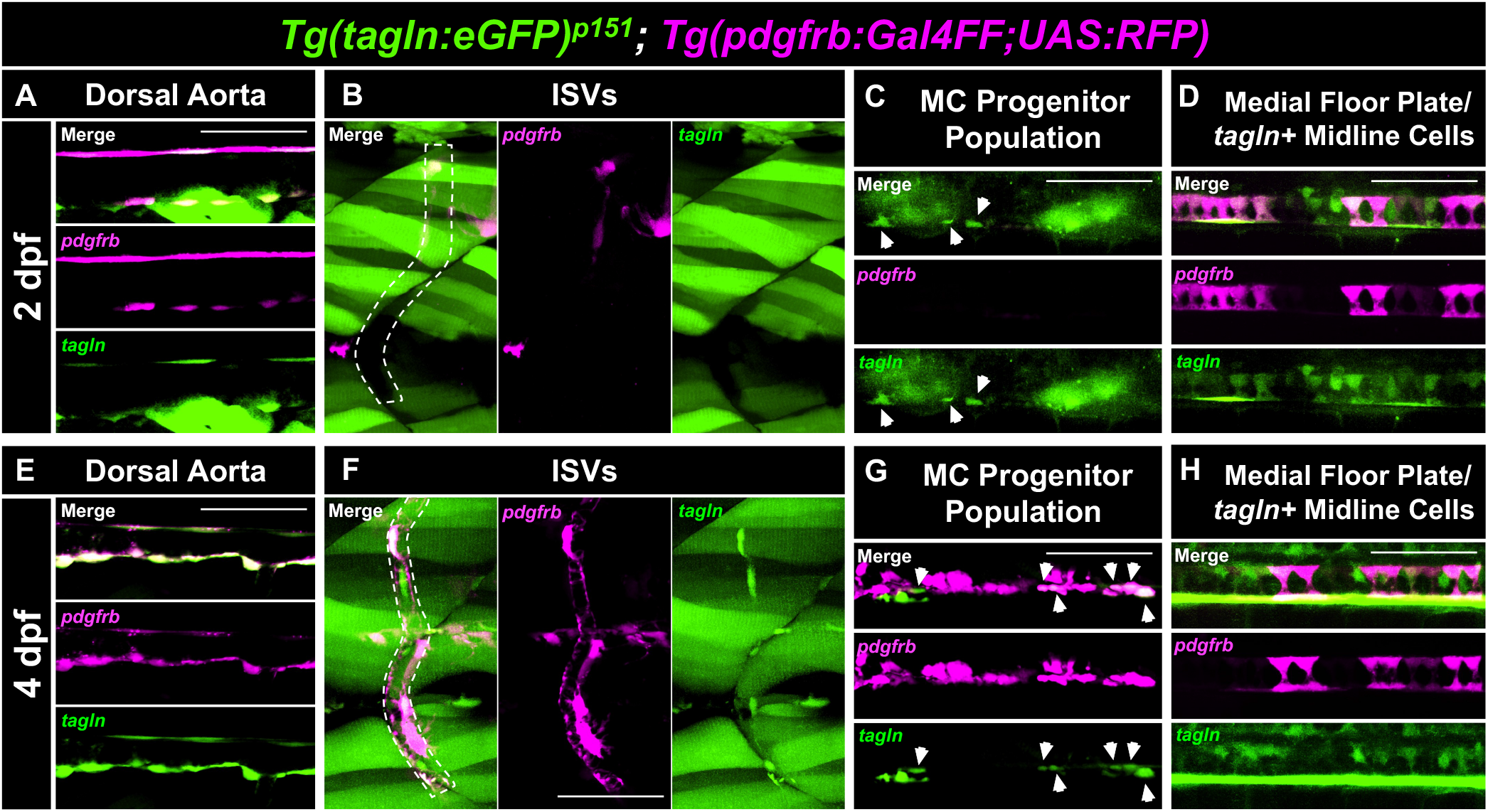
Visualizing *tagln* expression using a cytosolic-GFP *tagln* reporter line. Confocal images of *Tg(tagln:eGFP); Tg(pdgfrb:Gal4FF; UAS:RFP)* fish at 2 dpf (A-D) and 4 dpf (E-H). *tagln* expression was evident in the hypochord and vSMC populations of the dorsal aorta (A,E), though GFP signal was more easily detectable in vSMCs than the hypochord. GFP expression in the skeletal muscle decreased the efficiency and resolution of identifying *tagln*-positive cells in the trunk, though we were able to visualize *tagln*-high mural cells on the ISVs by 4 dpf (B,F). Despite high background, GFP signal was also detectable in the MC progenitor population and the medial floor plate cells (C,D,G,H). We observed intense GFP expression in the *tagln*-positive midline cells by 4 dpf (H). Thus, we were able to confirm *tagln* expression in these structures using an independent transgenic, the cytosolic-GFP *tagln* reporter. MC, mural cell. ISVs, intersegmental vessels. Scale bars: 50 µm.

## Movies 1, 2, 3

**Movie 1. A 3D view of *pdgfrb* and *tagln* expression in the hypochord between 1 to 4 dpf.** Movie of rotating 3D projections. Projections were made from confocal z-stacks of the hypochord in *Tg(tagln:NLS-EGFP); Tg(pdgfrb:Gal4FF; UAS:RFP)* zebrafish. *pdgfrb* expression decreased between 1-4 dpf, while cells of the hypochord elongated and decreased slightly in number, as evidenced by the change in shape and number of *tagln*-positive nuclei. Scale bar: 50 µm.

**Movie 2.** 3D views of MC progenitor cells migrating to the ISVs in *Tg(tagln:NLS-EGFP); Tg(pdgfrb:Gal4FF; UAS:RFP)* zebrafish at 3 dpf. Movie of two rotating 3D projections, side by side. Projections were made from confocal z-stacks of the MC progenitor population identified by *pdgfrb* and *tagln* expression. Arrows point to the migrating cells. ISVs are outlined by white dotted lines. Scale bar: 10 µm.

**Movie 3.** 3D views of MC progenitor cells migrating to the ISVs in *Tg(tagln:NLS-EGFP); Tg(Ola.Twist:GAL4; UAS:RFP)* zebrafish at 4 dpf. Movie of two rotating 3D projections, side by side. Projections were made from confocal z-stacks of the MC progenitor population identified by *twist1a* and *tagln* expression. Arrows point to the migrating cells. ISVs are outlined by white dotted lines. Scale bar: 10 µm.

